# Circadian regulation of lung repair and regeneration

**DOI:** 10.1101/2021.11.20.469376

**Authors:** Amruta Naik, Kaitlyn Forrest, Yasmine Issah, Utham Valekunja, Akhilesh B Reddy, Elizabeth Hennessy, Thomas S. Brooks, Apoorva Babu, Mike Morley, Gregory R. Grant, Garret A. FitzGerald, Amita Sehgal, G. Scott Worthen, David B. Frank, Edward E Morrisey, Shaon Sengupta

**Affiliations:** Department of Pediatrics, University of Pennsylvania Perelman School of Medicine; Institute of Translational Medicine and Therapeutics (ITMAT), University of Pennsylvania; Systems Pharmacology University of Pennsylvania Perelman School of Medicine; Department of Genetics, University of Pennsylvania Perelman School of Medicine; Department of Neuroscience, University of Pennsylvania Perelman School of Medicine; Lung Biology Institute, University of Pennsylvania Perelman School of Medicine

## Abstract

Optimal lung repair and regeneration is essential for recovery from viral infections such as that induced by influenza A virus (IAV). We have previously demonstrated that lung inflammation induced by IAV is under circadian control. However, it is not known if the circadian clock exerts its influence on lung repair and regenerative processes independent of acute inflammation from IAV. Here, we demonstrate for the first time that lung organoids have a functional clock as they mature and that the absence of an intact circadian clock impairs regenerative capacity. Using several models of circadian disruption, we show that with the absence of an intact clock lung proliferation is disrupted. Further, we find that the circadian clock acts through direct control of the Wnt/β-catenin pathway. We speculate, that adding the circadian dimension to the critical process of lung repair and regeneration will lead to novel therapies and improve outcomes. Finally, we use data from UK Biobank to demonstrate at the population level, the role of poor circadian rhythms in mediating negative outcomes following lung infection.

## INTRODUCTION

Circadian rhythms provide an anticipatory system by temporally compartmentalizing physiological processes and thus allow organisms to maintain homeostasis in the face of changes in their environment. At the molecular level, these cell-autonomous clocks comprise transcriptional-translational feedback loops of the core clock genes that drive the rhythmic fluctuations of many cellular processes^1^ through direct or indirect effects on downstream targets. Circadian profiling of the lung transcriptome reveals that most physiological pathways, including those involved in the regulation of lung injury and repair, are characterized by rhythmically expressed genes^2, 3^. Further, lung inflammation has been shown to be under circadian control in several injury models^2, 3^. We have previously demonstrated that the circadian clock determines acute mortality from Influenza A virus (IAV) infection in mice independent of viral burden^4^. Recovery from IAV infection is dependent not only on inflammation, but also the ability of the host to repair and regenerate injured lungs. However, the role of the circadian clock in lung repair and regeneration is not known.

Some studies have suggested a role for the circadian clock in tissue regeneration such as skin and intestines. This has been attributed to improved actin assembly in skin fibroblasts or through the regulation of cell proliferation in both skin and intestines^5–7^. However, both skin and intestines have very high turnover rates at baseline and are constantly proliferating and differentiating. The uninjured lung, on the other hand, is a quiescent organ in terms of proliferation. However, following acute inflammation induced by IAV infection, the lung enters a state of rapid proliferation^8^. Whether the circadian clock improves lung proliferation following IAV is not known. Our previous work supports the role of at least two pulmonary epithelial cells, contributing to the circadian control of acute outcomes from IAV, namely Sfptc^+^ Alveolar type 2 (AT2) cells and Scgb1a1^+^club cells^4, 9^. Acute mortality and lung injury was worse when the circadian clock was specifically disrupted in Sfptc^+^ or Scgb1a1^+^ cells^4, 9^. Interestingly, both these cell types are central to lung repair and regeneration following IAV infection. Thus, we hypothesized that circadian clock improves long term recovery from IAV through lung repair and regenerative pathways and, this effect is at least partly independent of the clock-gated inflammation.

To determine the extent to which the circadian clock contributes to lung regeneration independent of clock-gated inflammation, we employ lung organoid assays. We demonstrate for the first time, that lung organoids have their own intrinsic clock, and that disruption of the circadian clock impairs lung regenerative responses. Using a combination of tissue specific and constitutive *Bmal1^-/-^* and constitutive *Cry*1^-/-^Cry2^-/-^models, we confirm the role of circadian control in directing lung repair and regeneration. Further, we use epithelial cell-specific clock mutants and single cell RNA sequencing to demonstrate that the clock aids lung repair and regeneration by controlling proliferation during recovery from IAV infection. Specifically, we identify circadian control of the wingless related-integration site (Wnt) /β-catenin pathway as a mechanism for clock-gated cell proliferation.

## RESULTS

### Temporal gating of alveolar damage on day 30 post-recovery is controlled by the circadian clock

We have previously shown that mice infected at dusk (ZT11) had a 3-fold higher mortality than littermates and cagemates infected at dawn (ZT23) with lethal doses of the virus. [By circadian nomenclature, ZT0 refers to the time when lights turn on in a 12hr light-dark cycle. Thus, ZT11 or dusk refers to the time just before the onset of the rest phase and ZT23 refers to the time just before the onset of active phase in mice who are nocturnal]. To determine whether the time of day at initial infection affects lung repair later, we infected C57bl6J mice with IAV (H1N1; PR8; sub-lethal dose) at ZT11or ZT23 and harvested lungs 30 days post-infection (Fig 1A). No significant mortality was noted at this dose in either group. Interestingly, we found that even one month after the infection, mice infected at ZT11 had significantly more lung damage and alveolar destruction than the group infected at ZT23 using a previously validated scoring^10^ (Zone 3 or severe injury: 15.87%- in ZT23 versus 27.37% in ZT11, , p<0.007; Zone 4 or complete alveolar destruction: 10.04% in ZT23 versus 18.52% in ZT11 group, p<0.02, Mann-Whitney test), suggesting that the temporal protection from circadian rhythmicity persists well beyond the acute phase of inflammation (Fig 1B-C). These data raise the possibility that the clock modifies lung repair and regeneration that typically follow acute inflammation. However, in the experimental paradigm used above, the day 30 histology reflects both the role of the clock in initial inflammation and any potential contribution to lung repair and regeneration. Thus, to study the role of the circadian clock in lung regeneration independent of inflammatory influences, we used 3-D organotypic assays from the lung epithelium.

**Figure 1.**
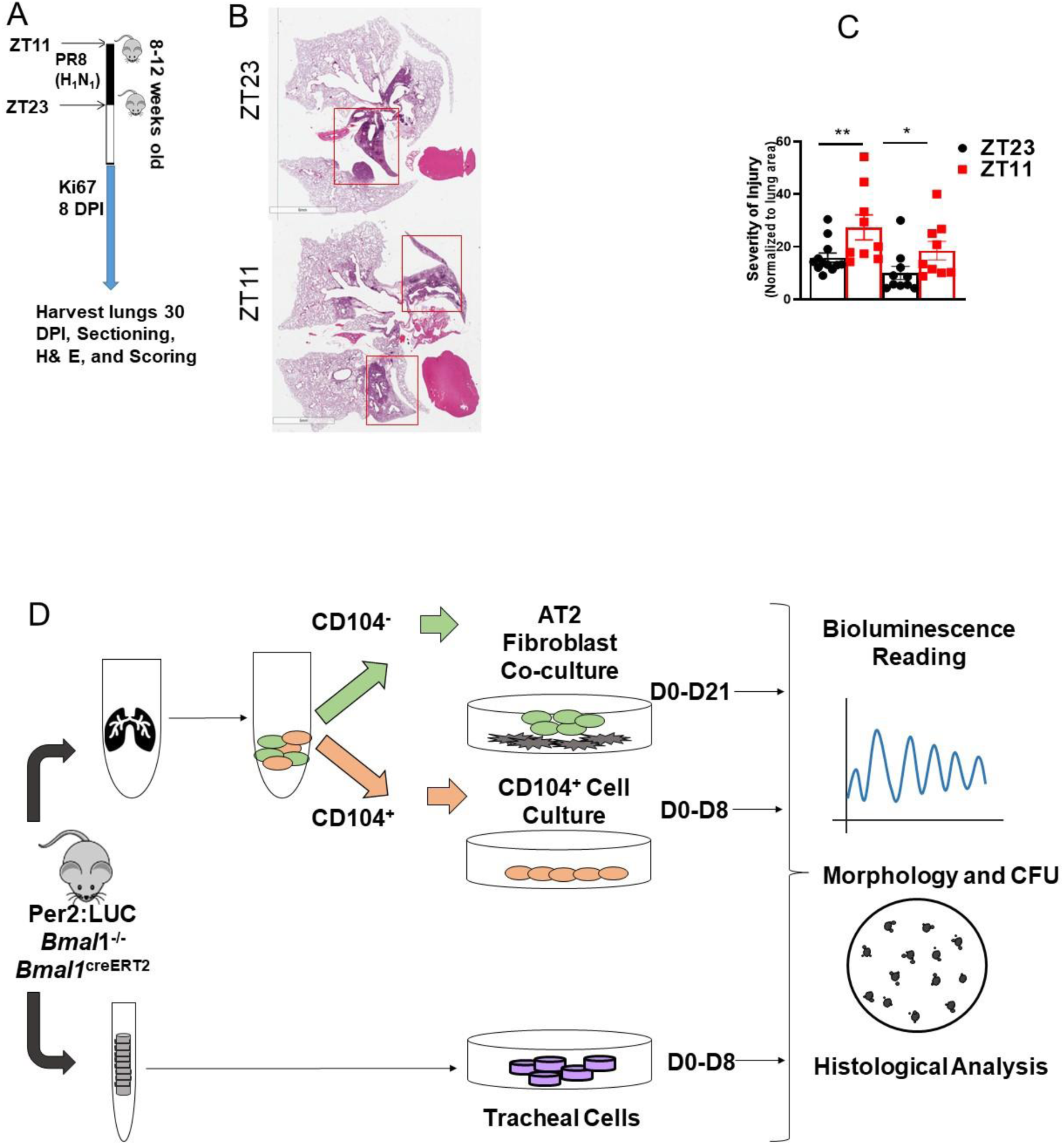
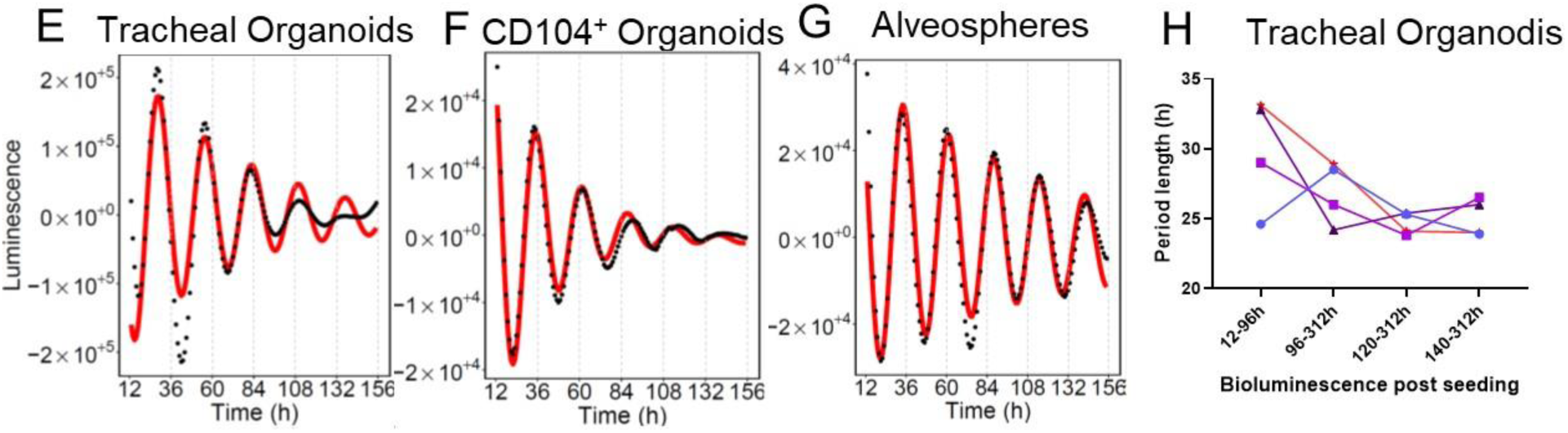

### Organoids from different levels of the respiratory tract display robust circadian rhythmicity

However, before examining the impact of circadian rhythms on regenerative response, we wanted to determine if the organoids maintained an active clock. While many studies have demonstrated the presence of a cell autonomous clock active in *in vitro* cultures, evidence of the same in organotypic cultures is limited, mainly restricted to rapidly proliferating, intestinal organoids or metabolically active, β (islet) cell organoids^6, 7, 11^. To check for the presence of an intact circadian clock in organoids, we used the *mPer2^luc^* mice^12^. Lung organoids were grown from different levels of the respiratory epithelium, namely alveolar, small airway and tracheal regions, of the *mPer2^luc^* (Fig 1D).

Lung organoids from these regions exhibited robust diurnal oscillations, even in the absence of synchronizing agents, for more than 108 h for tracheal organoids (Fig 1E, Table 2, period: 23.8h, phase: 0h), > 108 h for CD104^+^ organoids from smaller airways (Fig 1F, Table 2, period: 26.2 h, phase: 9.3h), and >204 h for alveolar organoids (Fig 1G, Table 2, period: 27.2h, phase: 6.5).

Addition of the synchronizing agents dexamethasone (10 µM) or forskolin (100nm) extended the duration of observed circadian rhythmicity of all the organoids. (S. Fig 2 A-F, Table 2). Of note, we observed that circadian rhythmicity develops as a function of maturity. Tracheal organoids mature by day 7 in culture and the bioluminescence data in Fig 1E were recorded after 7 days following seeding. However, when the recording was started on day 4 after seeding, the period length was substantially variable and longer. As the tracheal organoids matured, the period length decreased to a “24 hr” time period, consistent with circadian rhythmicity and was noted to be much less variable (Fig 1H). These results confirm the existence of a cell intrinsic clocks in lungs organoids and further support their use in studying the effect of the circadian rhythms on lung regeneration. Moreover, it suggests that the process of maturation/regeneration in organotypic cultures is paralleled by consolidation of circadian rhythms.

### Loss of *Bmal1* reduces the regenerative potential of tracheal and distal lung basal cell organoids

Next, we examined the role of the circadian clock in modulating the regenerative capacity of epithelial cells capable of self-renewal and/or differentiation from all levels of the respiratory tree (Fig 1D). Only cells capable of regeneration contribute to the growth of these self-organizing structures. These include alveolar type 2 cells (AT2), basal cells and club cells^13–16^. Organoids grown from alveolar regions reflect the regenerative capacity of AT2 cell. As there are no established unique surface markers for sorting live club cells, we used the following strategy. Organoids grown from the CD104^+^ epithelial cells reflect the regenerative capacity of a combination of club cells and basal cells from smaller airways including a population of CD104^+^ lineage-negative epithelial progenitor^17^ and SOX2^+^ Lin^-^ cells^18^, both known for their role in lung repair. Tracheal organoids reflect the regenerative capacity of club cells and basal cells but from trachea/larger airways. For the experiments below, all cells were harvested from mice with global *Bmal1* deficiency.

We found that embryonic loss of *Bmal1* resulted in a 54% decrease in colony forming efficiency (Fig. 2A-B, E; p<0.0005, unpaired t test with Welch’s correction, CFE) in tracheal organoids. Similarly, loss of *Bmal1* also decreased the regenerative capacity of CD104^+^ organoids by 24% (Fig 2G-H, K; p<0.01, unpaired t test, Welch’s correction) in comparison to organoids derived from the *Bmal1* sufficient CD104^+^ lung cells. Although *Bmal1* is the only non-redundant core clock gene, whose deletion alone is sufficient to produce a dramatic, arrhythmia, the global phenotype is exaggerated by its non-circadian functions^19^. If the regenerative loss seen in the embryonic knockout^-^ model is truly secondary to its circadian role, we argued that this phenotype should be preserved in a postnatal model wherein the clock has been disrupted in adulthood (∼8 weeks of life).

**Figure 2.**
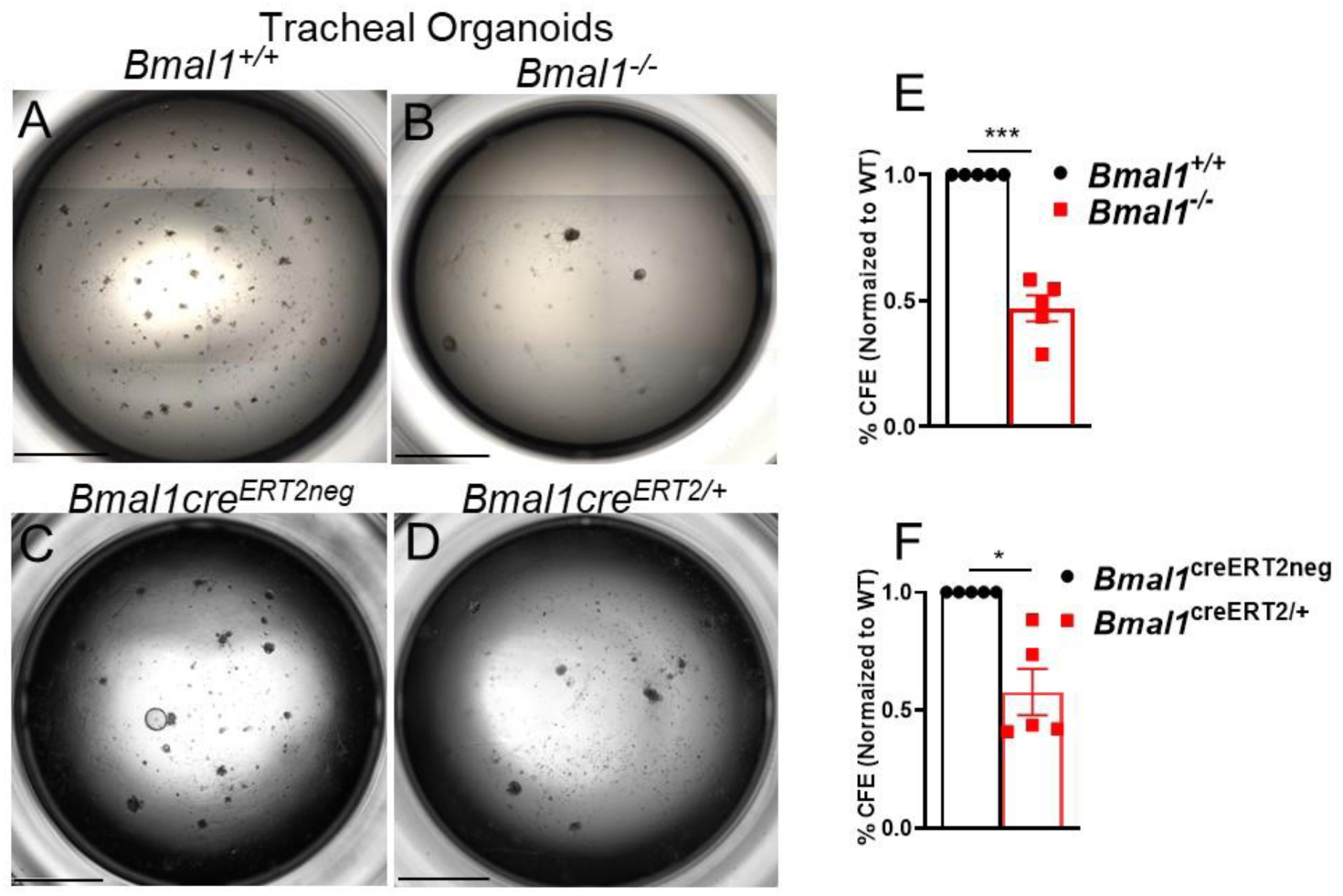
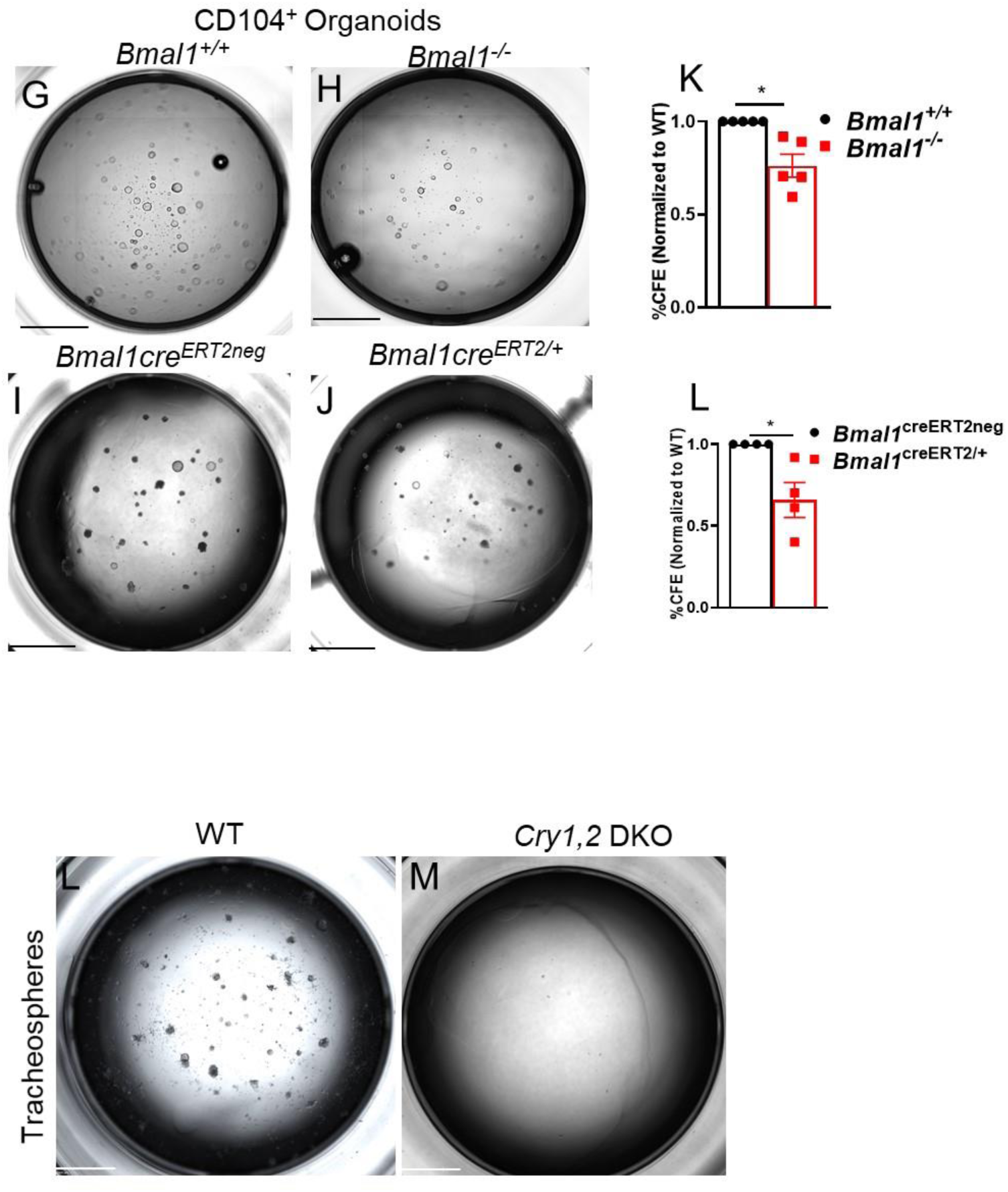
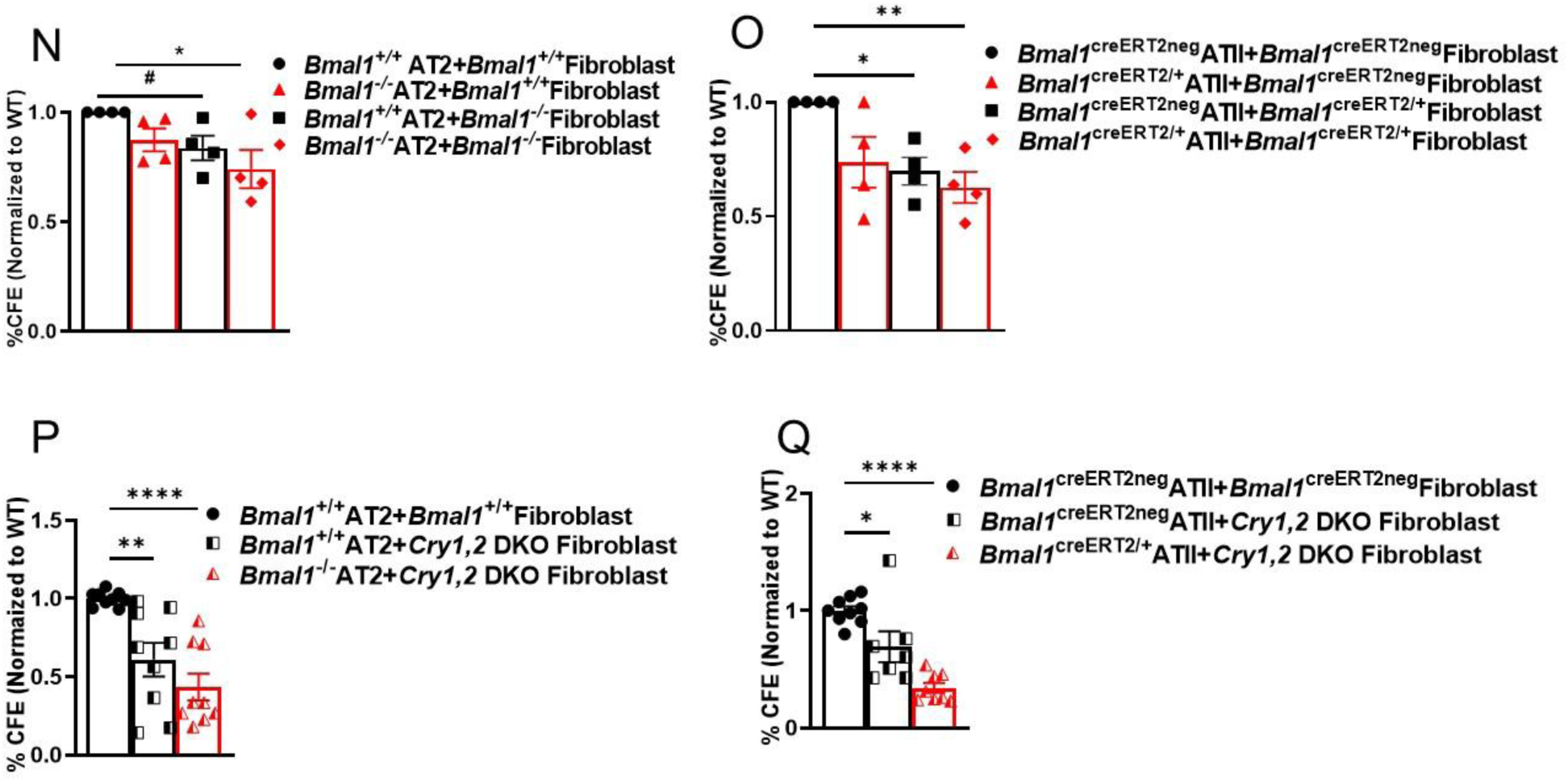

Indeed, we found that even postnatal loss of *Bmal1* (*Bmal1*^creERT2/+^) decreased the regenerative capacity of tracheal organoids by 43% (Fig. 2C-D, F; p<0.01, unpaired t test, Welch’s correction) and of CD104^+^ organoids by 35% (Fig 2I-J, L; p<0.04, unpaired t test, Welch’s correction) as compared to *Bmal1*^creERT2neg^derived organoids.

To test if this loss of regenerative capacity was *Bmal1* specific or broadly recapitulated in other models of clock disruption, we used lung cells from *Cry*1^-/-^*Cry*2^-/-^ (*Cry1, 2* DKO) mice. Tracheal organoids derived from *Cry1, 2* DKO mice show drastically reduced regenerative capacity, with hardly any organoids seen in the mutant (Fig 2L-M). Taken together these findings confirm that an intact circadian clock improves lung recovery through enhanced regeneration, independent of any influence from clock-gated inflammation. To understand whether the poor regenerative capacity due to clock disruption is specific to epithelial cells in larger airways (tracheal cells and bronchial cells) or globally relevant to all cell types, we investigated the effect of *Bmal1* silencing on alveolar cells.

### Clock disruption reduces the regenerative capacity of the alveolar cells

To test the effect of clock disruption on the regenerative ability of AT2 cells, we cultured CD45^-^ CD31^-^EpCAM^+^CD104^-^cells comprising both AT2 and AT1 cells (S Fig 1A) in organotypic assay. As described in the previous section, we used both the global embryonic and postnatal inducible *Bmal1*^-/-^ models. Lung fibroblasts from these mice and their respective WT littermates were used as feeder cells, to support AT2 cells self-renewal^13, 20^. To distinguish between the contribution of *Bmal1* in the mesenchyme versus the epithelium to the overall regenerative capacity of the lung alveoli, we used the following four experimental combinations (1) *Bmal1*^-/-^AT2 with *Bmal1*^-/-^ fibroblasts (2) *Bmal1*^-/-^AT2 with *Bmal1*^+/+^ fibroblasts (3) *Bmal1*^+/+^AT2 with *Bmal1*^-/-^ fibroblasts, and (4) *Bmal1*^+/+^ATs with *Bmal1*^+/+^ fibroblasts from both embryonic and inducible models(S. Fig 3A- H).

There was a 13% decline in regenerative capacity with embryonic deletion of *Bmal1* in AT2 cells supported by WT fibroblasts (Fig 2N, S. Fig 3B), while there was a 17% reduction when *Bmal1* was deleted embryonically in fibroblasts but not in AT2 cells (S. Fig 3C, Fig 2N, p=0.06, unpaired t test, Welch’s correction). Regenerative ability decreases compounded (26%) when both the AT2 cells and fibroblasts were deficient in *Bmal1* (S. Fig 3D, Fig. 2N, p<0.05 ordinary one-way ANOVA).

Interestingly, the loss of regenerative potential was more marked for alveolar organoids derived from cells following postnatal deletion of *Bmal1* than from those following the embryonic loss of *Bmal1.* There was a 27% reduction in regenerative capacity in *Bmal1*^creERT2/+^AT2s co-cultured with WT feeder cells (S. Fig 3F, Fig 2O), and 31% decrease with WT AT2 and *Bmal1*^creERT2/+^feeder fibroblasts (S. Fig 3G, Fig 2O p<0.01, Ordinary one-way ANOVA). The ability of *Bmal1*^creERT2/+^AT2s to regenerate when co-cultured with *Bmal1*^creERT2/+^fibroblasts was further reduced by 38% (S. Fig 3H, Fig 2O, p<0.005, one-way ordinary ANOVA). This effect was not limited to a *Bmal1* specific effect, and similar deficiency was noticed when *Cry1, 2* DKO cells were used. The regenerative capacity of alveolar organoid assays was decreased by 44% and 57% for *Bmal1*^+/+^ alveolar organoids (p<0.003, one-way ANOVA), *Bmal1*^-/-^ (Fig. 2P, p<0.0001, one-way ANOVA) when supported by *Cry1, 2 DKO* fibroblasts. The colony forming efficiency of *Bmal1*^creERT2/+^AT2s was furthermore reduced by 66% when co-cultured with *Cry1, 2 DKO* fibroblasts (S. Fig 3K, Fig 3Q p<0.0001, one-way ANOVA). Overall, our data are consistent with the circadian clock aiding lung regeneration via roles in both the alveolar epithelial and mesenchymal cells. When comparing the two models of *Bmal1* deletion, we noted that the organoids derived from the embryonic *Bmal1* deletion model, demonstrated a rather dysplastic appearance across all three types of organoids (tracheal, CD104^+^ and CD104^-^ , S. Fig 4E-H). This was not present in organoids derived from the inducible model (S. Fig 4L-O). Overall, the sizes of the organoids were comparable. In summary, while subtle differences in organoid appearance may be present in the two *Bmal1* deletion models, they remain comparable at least in regenerative efficiency.

**Figure 3.**
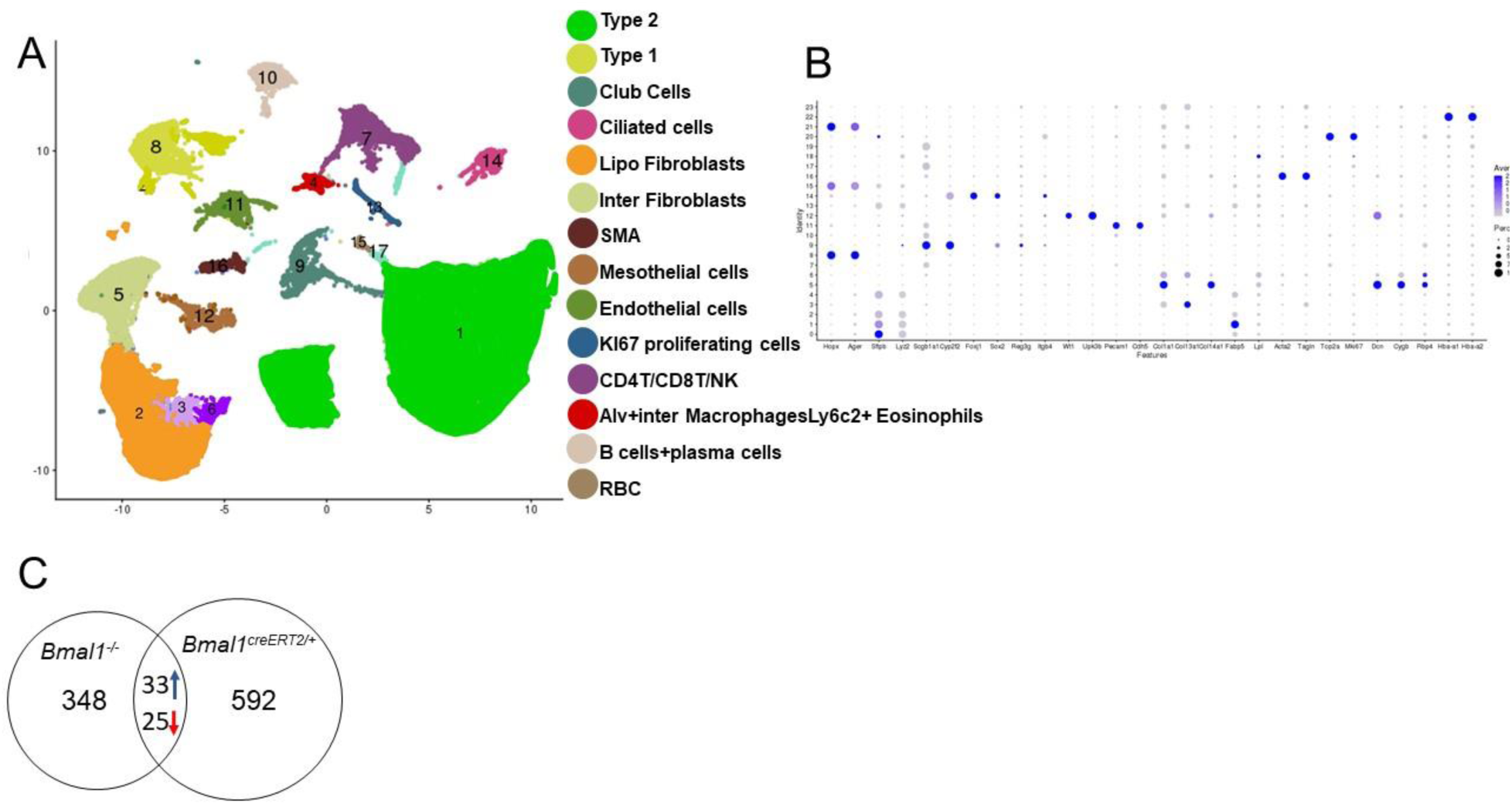
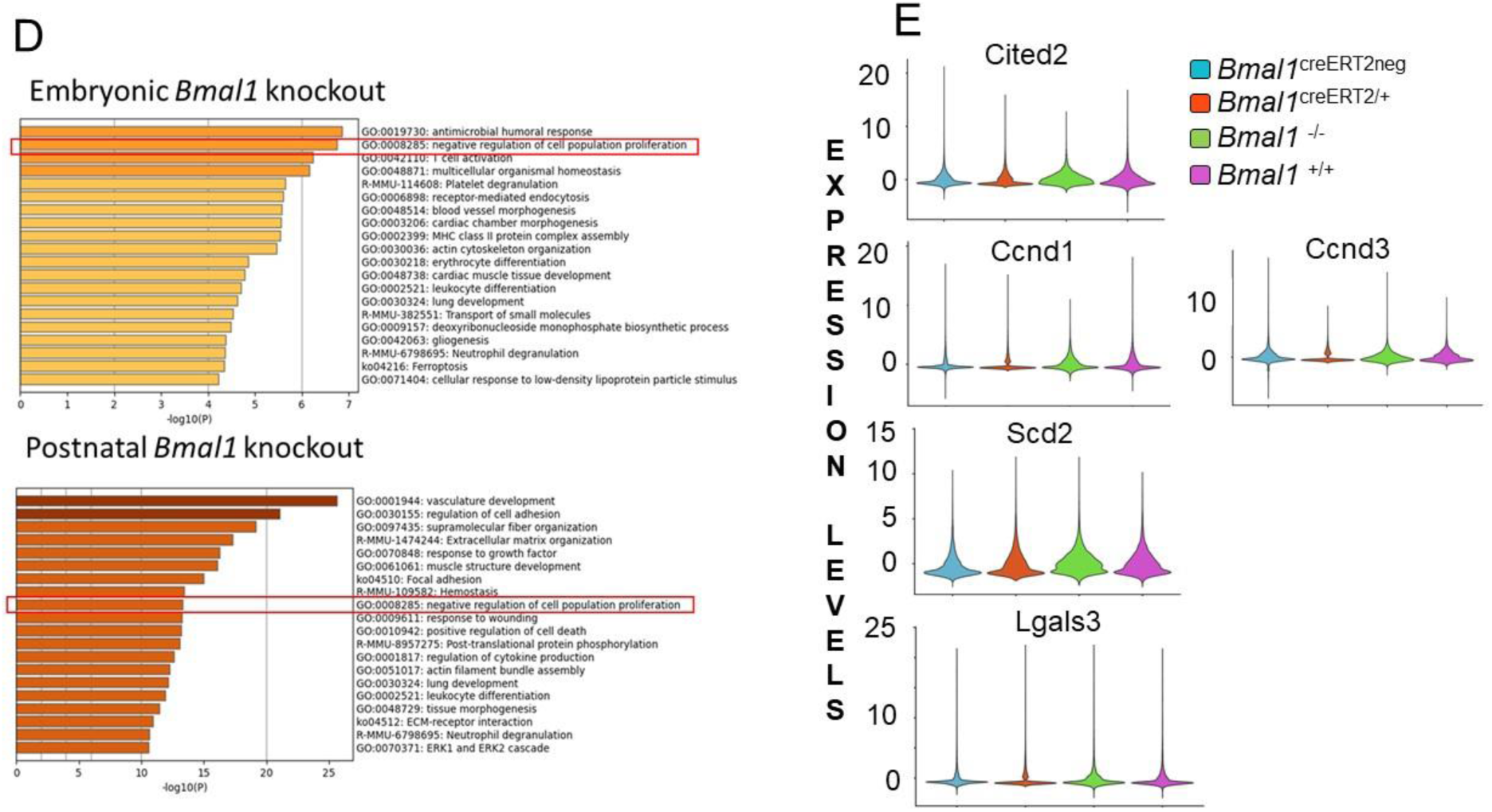

### Single-cell transcriptomic analysis of embryonic and postnatal knockouts reveals differences in the Wnt3a pathway

To elucidate the mechanistic basis of the regenerative deficiency seen with clock disruption we undertook an unbiased approach with single-cell RNA sequencing (scRNA-seq) of epithelial and mesenchymal cells from *Bmal1*^-/-^, *Bmal1*^creERT2/+^, and their respective WT littermates (S. Fig 1B). Because of the similarity seen between the two knockout models, we integrated the data from both knockout models with scRNA-seq data obtained from their wild type littermate controls. Projection of transcriptomic variations of individual cells by uniform manifold approximation and projection (UMAP) showed 17 clusters (Fig 3A). Clusters were annotated based on the differential gene expression analysis of cell-type specific known marker genes with highly different levels between clusters^21, 22^(https://research.cchmc.org/pbge/lunggens/) (Fig 3B). A total of 406 genes in the embryonic knockout model, and 650 genes in the postnatal knockout model were differentially expressed (source file 3). Common to both models of *Bmal1* deletion, in comparing the *Bmal1^-/-^* cells vs WT, 33 genes were upregulated, and 25 genes were downregulated (Fig.3C, source file 3, adjusted p value<0.05, Wilcox method). We examined the top twenty GO categories and discovered that genes involved in negative regulation of cell proliferation and lung development were highly enriched in both knockout models (Fig 3D). Several genes implicated in regulating proliferation were identified in our scRNA data. Cited2 ((Cbp/p300 Interacting Transactivator with Glu/Asp Rich Carboxy-Terminal Domain 2), a negative regulator of Hif1-ᾳ whose gene product Cbp/p300 interacts with β-catenin, and is involved in the transcriptional regulation of Wnt signaling was also downregulated^23^. Cyclin D promotes cell proliferation as a regulatory partner for CDK4 or CDK6 and its homologues Ccnd1 and Ccnd3 were dowregulated in *Bmal1^-/-^* lungs. Ccnd1 is a transcriptional target of Wnt/ β-catenin canonical pathway^24, 25^. Another Wnt/β-catenin target gene, stearoyl-CoA desaturases2 (*Scd2*) was found to be upregulated in both knockout models^26^. Lgals3 was amongst one of the upregulated genes, and its product galectin-3 is a key regulator in the Wnt /β-catenin signaling pathway through its interactions with axin-2, GSK-3β and β-catenin^27–29^. Wnt/β-catenin signaling pathway plays an instrumental role in stem cell self-renewal. Ccnd1 and β-catenin are two of the many clock-controlled cell cycle regulators. Activated canonical Wnt signaling has been shown to be involved in differentiation of lung epithelial stem/ progenitor cells^10, 20, 30, 31^. We identified targets of Wnt/ β-catenin pathway those show rhythmic oscillations in lung tissue (Table 3) along with Cbr2, Ctsl, Pdgfrᾳ the genes those were differentially expressed in both KO models. Hes1 a direct target of the Notch pathway, was downregulated in *Bmal1*^-/-^ lungs. Given the importance of the Notch and Wnt pathways in mediating lung proliferation and differentiation and their contribution to recovery from IAV, we next investigated if the circadian clock was relevant to this process.

### Circadian disruption impairs cell proliferation in the lungs infected with influenza

To determine whether clock-gated lung repair is associated with selective deficiencies in the cellular composition of the lung epithelium, we stained the lungs for AT1 and AT2 markers from C57bl6J 30 day post IAV recovery model explained in Fig 1A. No differences were observed in the absolute number of Sftpc^+^ AT2 cells or Pdpn^+^ AT1 fluorescence intensity in either ZTs (Fig 4A-B). We next investigated if the circadian clock contributes to lung repair through the proliferation that follows acute injury from IAV. To do so, we enumerated Ki67^+^cells in lungs of IAV infected *Cry1, 2 DKO* animals. Notably, only 9.6% cells retained their proliferative ability in the *Cry1, 2 DKO* mice, affirming the role of the clock in cell proliferation post lung injury (Fig 4C-E, p<0.0129, unpaired t test, Welch’s correction).

**Figure 4.**
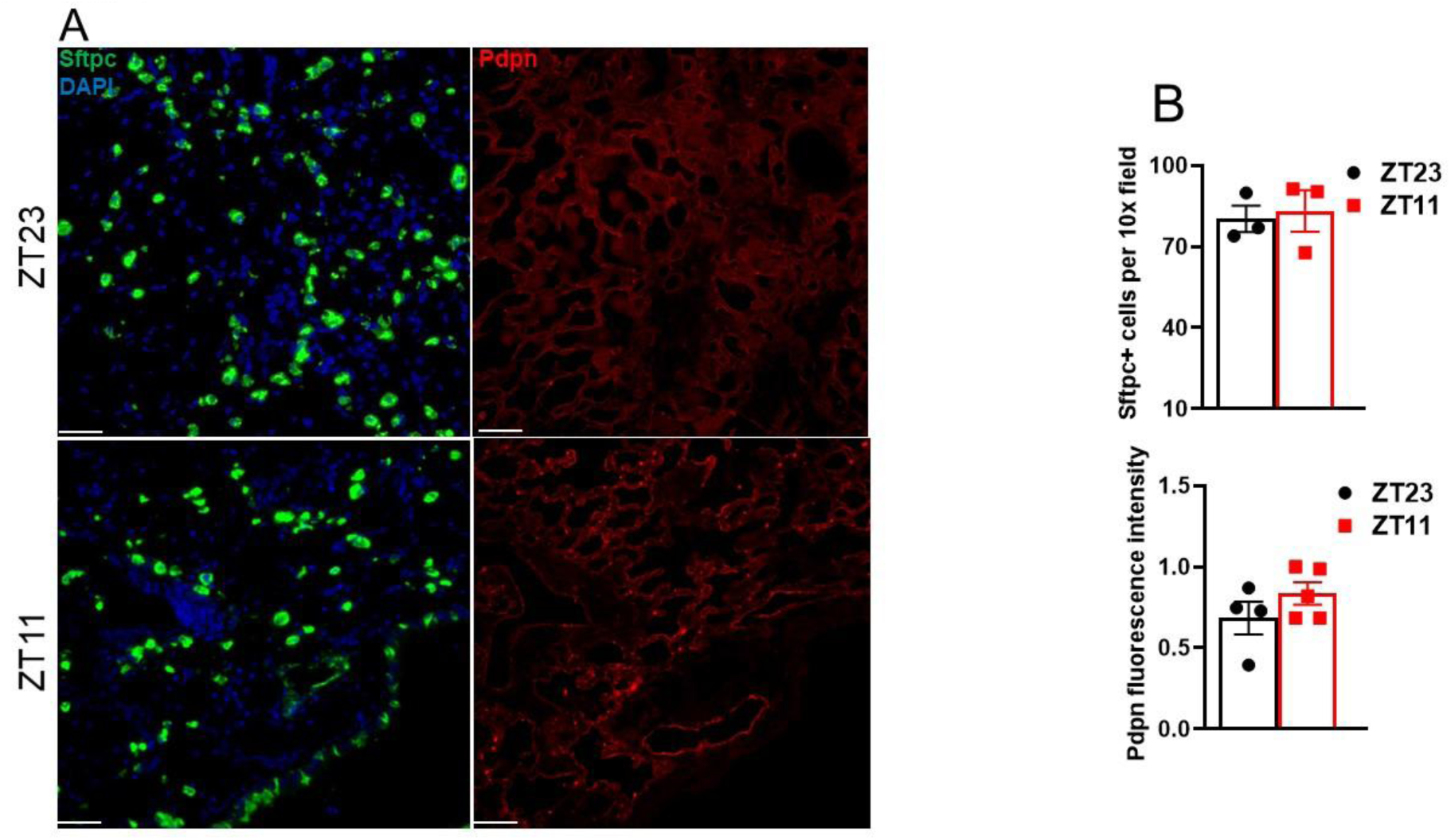
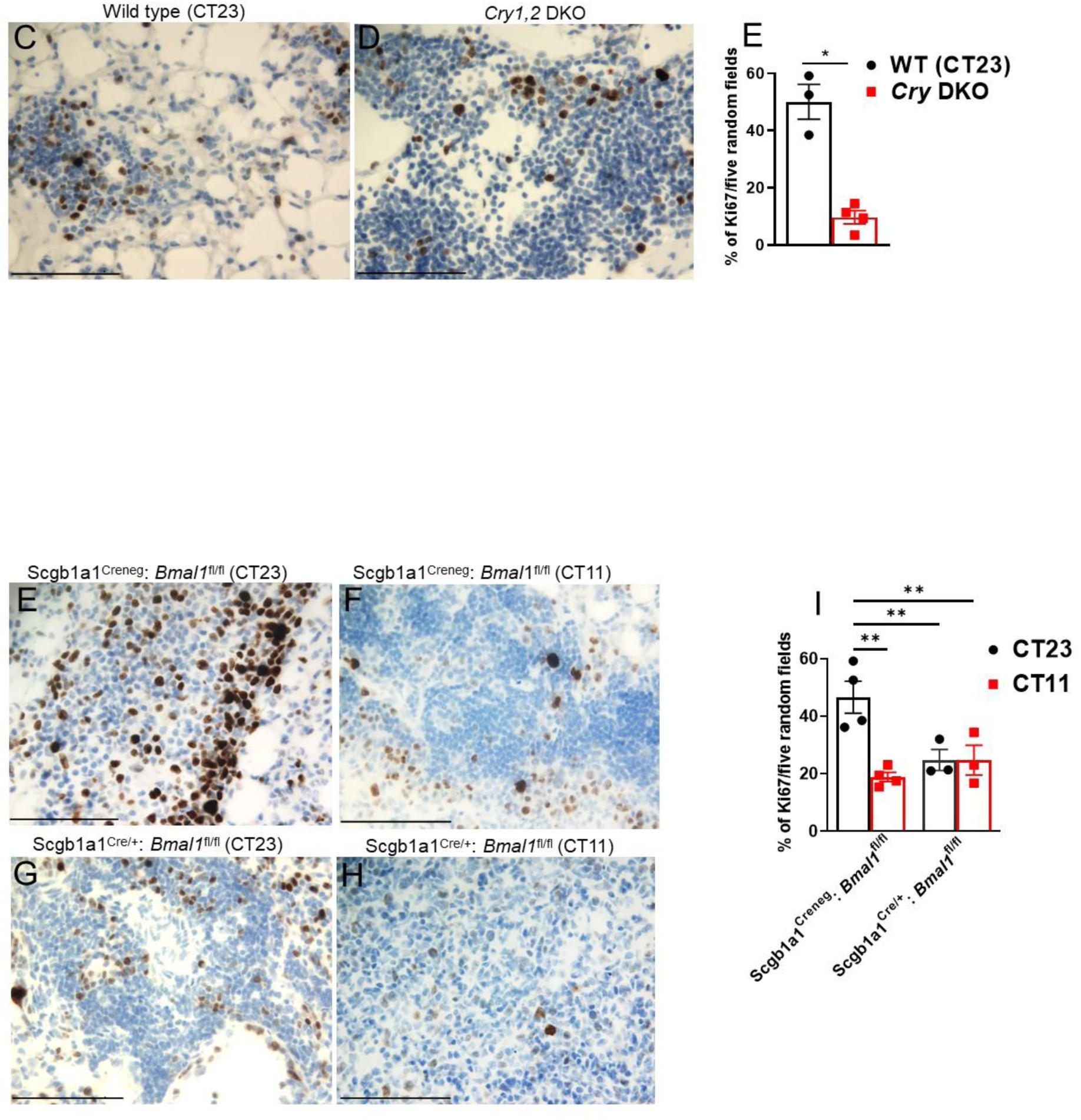
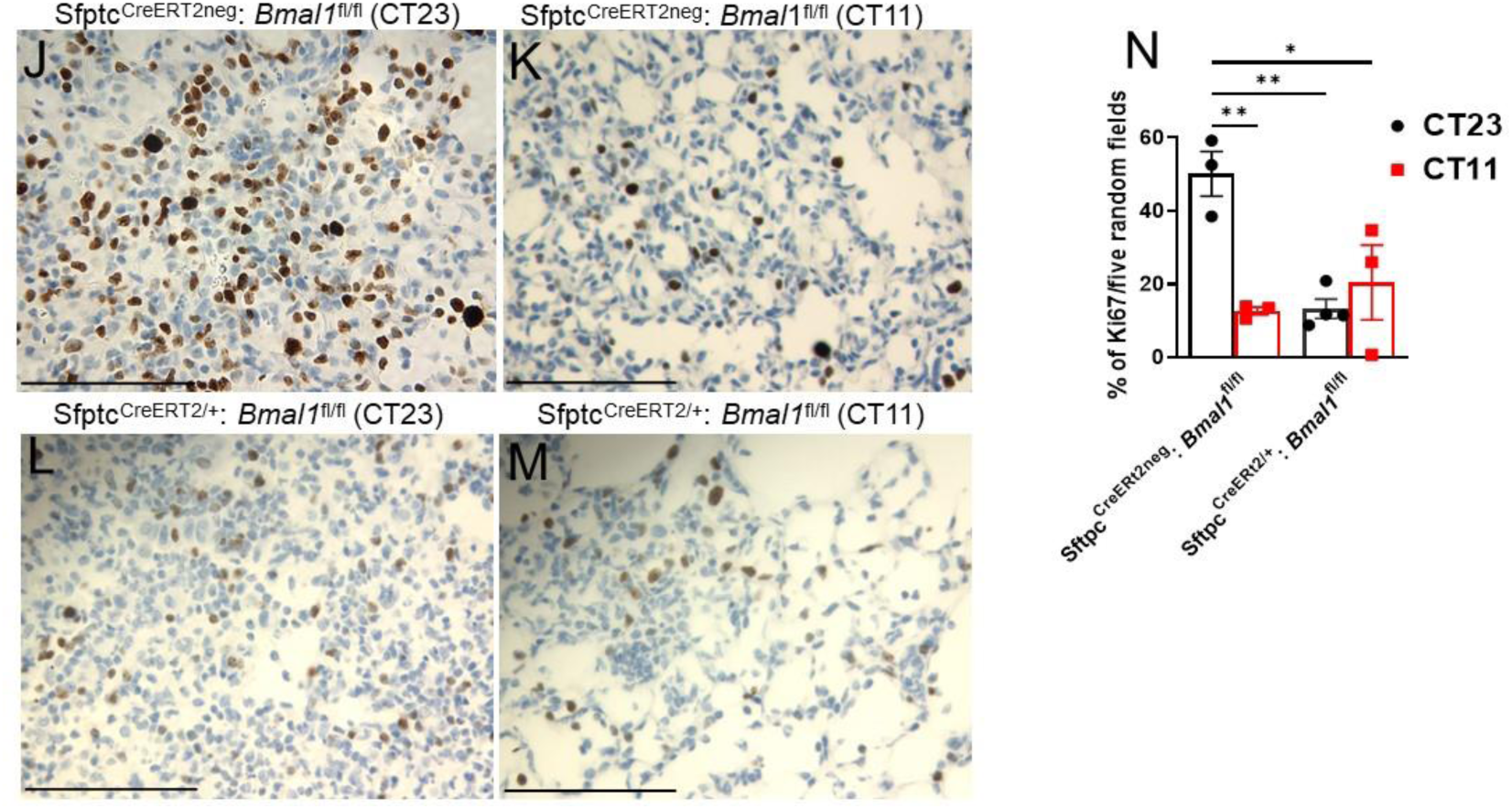

These data are consistent with the possibility that the worse histopathological outcomes 30 days post-infection in mice infected at ZT11 rather than ZT23, is likely to due to lower proliferation rather than transdifferentiation.

Having demonstrated that tracheal organoids from both *Bmal1* knockout models exhibit reduced regenerative capacity, we next investigated whether the club cell clock facilitates cellular proliferation in the aftermath of IAV *in vivo (Bmal1* was selectively deleted in club cells at embryonic stage). In *Scgb1a1^Cre/+^: Bmal1^fl/fl^* both cre+ groups had lower proliferation than the *Scgb1a1^Creneg^:Bmal1^fl/fl^*mice infected at CT23 (Fig 4G-H). However, the temporal difference in cell proliferation was still maintained in *Scgb1a1^Creneg^:Bmal1^fl/fl^*mice where mice infected at CT11 showed 2.5 fold decrease in proliferation than those infected at CT23 (Fig 4E-F, I; p<0.003, Ordinary one-way ANOVA).

Since alveolar repair is dependent on proliferation of AT2 cells, we infected Sftpc^Cre-ERT2/+^: *Bmal1*^fl/fl^ (wherein *Bmal1* was selectively deleted in AT2 cells), along with their cre^neg^littermates at either CT11 or CT23 and quantified Ki67+ and Sftpc^+^ Ki67+ cells on lungs harvested 8 days post IAV infection. Significantly fewer proliferating cells (12%) were observed in animals infected at CT11 versus CT23 (50%) in Sftpc^CreERT2neg^: *Bmal1*^fl/fl^ mice (Fig 4J-K, N**p<0.004, *<0.05, Ordinary one-way ANOVA). Conversely, this time-of-day difference was lost in Sftpc^Cre-ERT2/+^: *Bmal1*^fl/fl^littermates, suggesting that loss of *Bmal1* in Sftpc^+^ cells alter the number of proliferating cells (Fig 4L-M). Similarly, significantly fewer proliferating AT2 cells were observed in SftpcCre-ERT2/^+^: *Bmal1*^fl/fl^ than their cre negative littermates (S. Fig 3M). Collectively, our data shows that the circadian clock modulates regenerative capacity by impairing proliferation in the aftermath of IAV.

### *Bmal1* directly binds to E-box motif in Wnt3a promoter

Although the connection between the circadian clock and the Wnt/β-catenin pathways has not been investigated before, in adipose tissue, wnt activation was reduced specifically in in *Bmal1^-/-^*mice which in turn suppressed adipogenesis^32^. Another study has demonstrated that the circadian clock synchronizes the timing of cell division via rhythmic secretion of Wnt3a in intestinal organoids^6^. Based on our results from the sc-seq and lung proliferation, we investigated whether core clock gene can direct regulate the transcriptional activity of the Wnt/β-catenin pathway. Canonical Wnt signaling involves β-catenin stabilization and accumulation in the cytosol. Thereafter, translocates into the nucleus and activates downstream genes. First, we analyzed β- catenin abundance in the cytosol as a measure of Wnt activity in naïve lungs. Cytosolic β-catenin expression was significantly lower in the lung protein extracts of the *Bmal1*^creERT2/+^than in their *Bmal1*^creERT2neg^littermates suggesting that *Bmal1*, may directly target the Wnt pathways (Fig 5A-B). In turn, this downregulation at baseline could result in impaired activation of downstream Wnt-signaling and account for the loss of regenerative capacity seen in our models of clock disruption.

**Figure 5.**
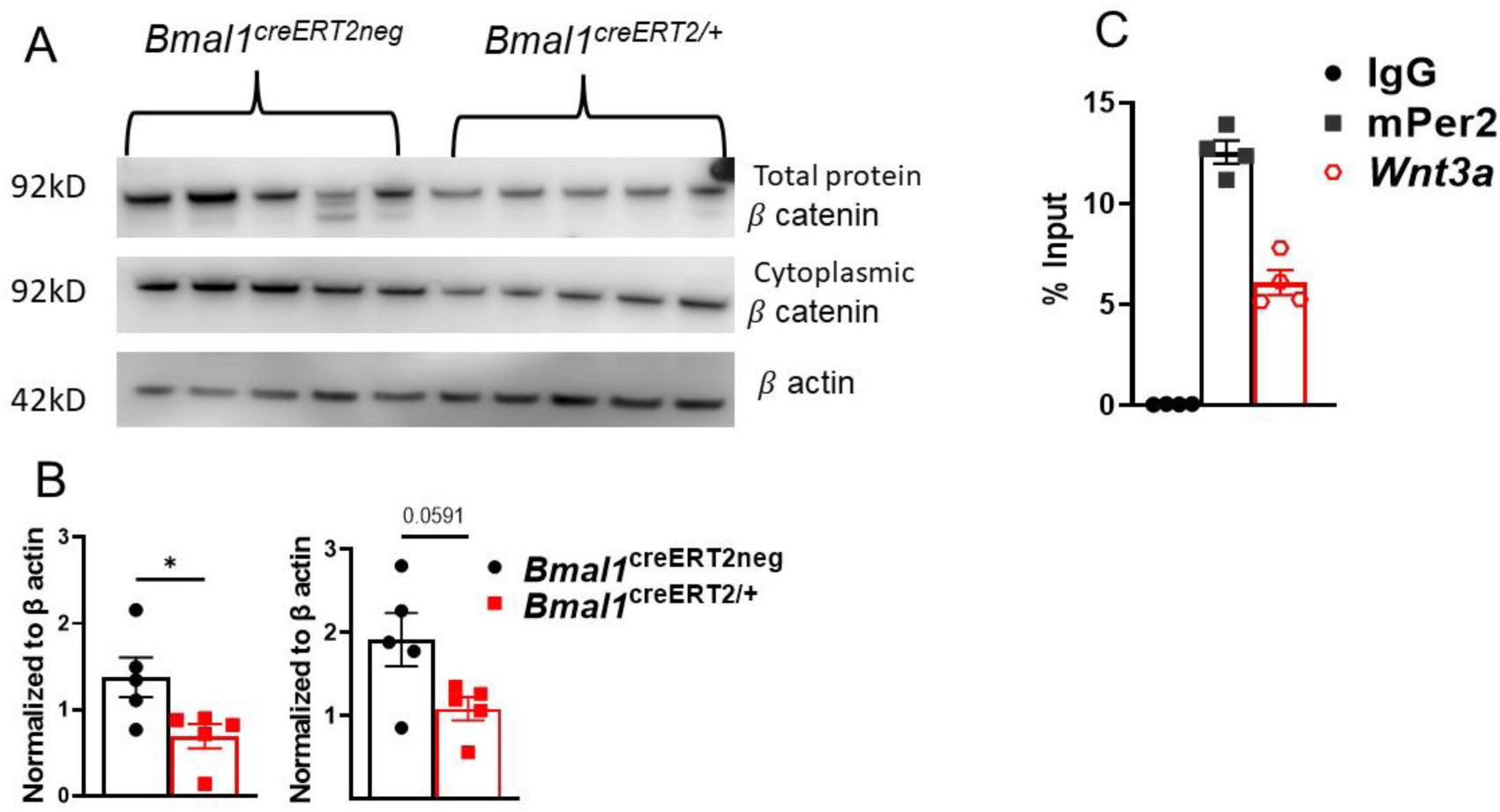
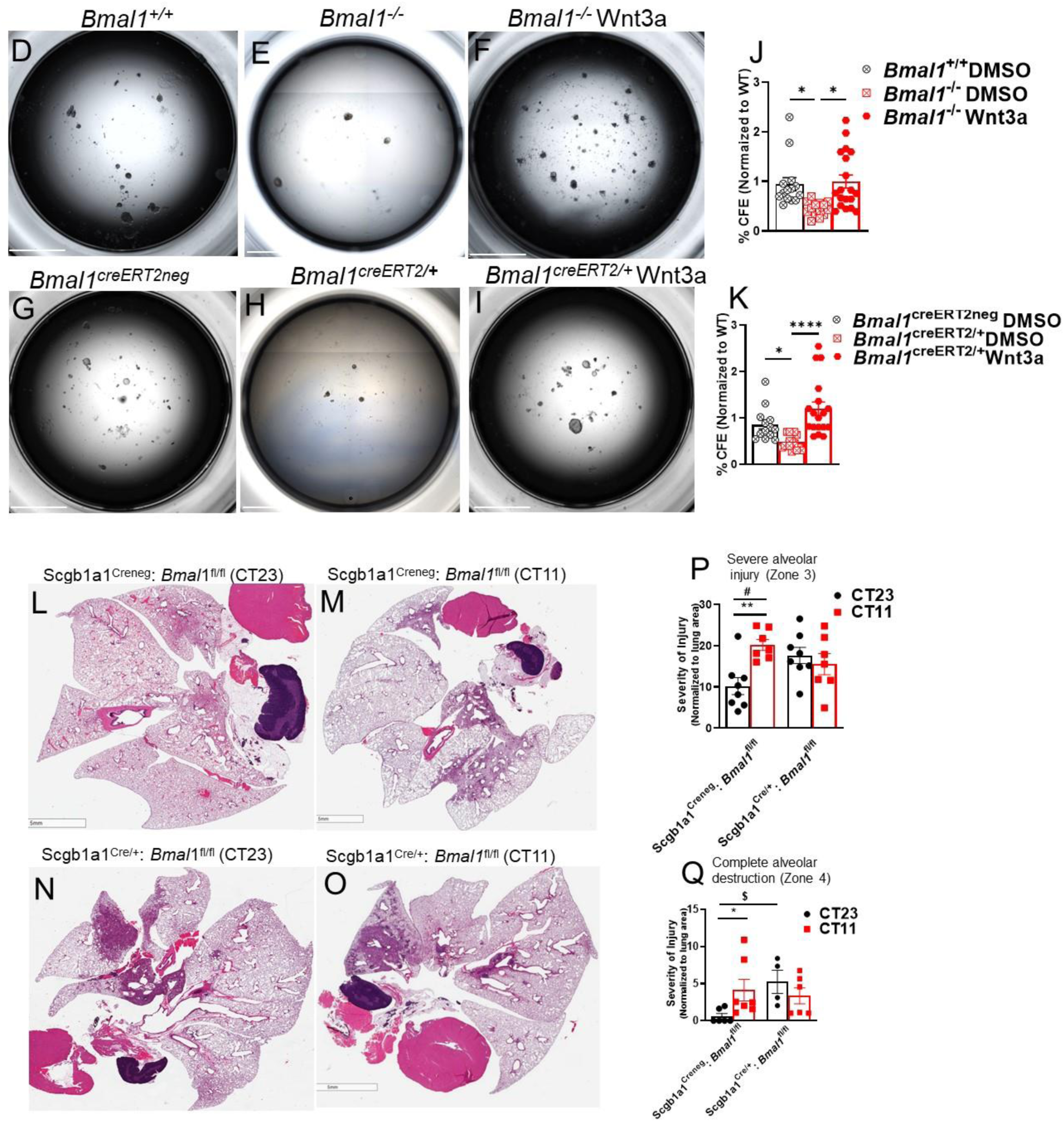
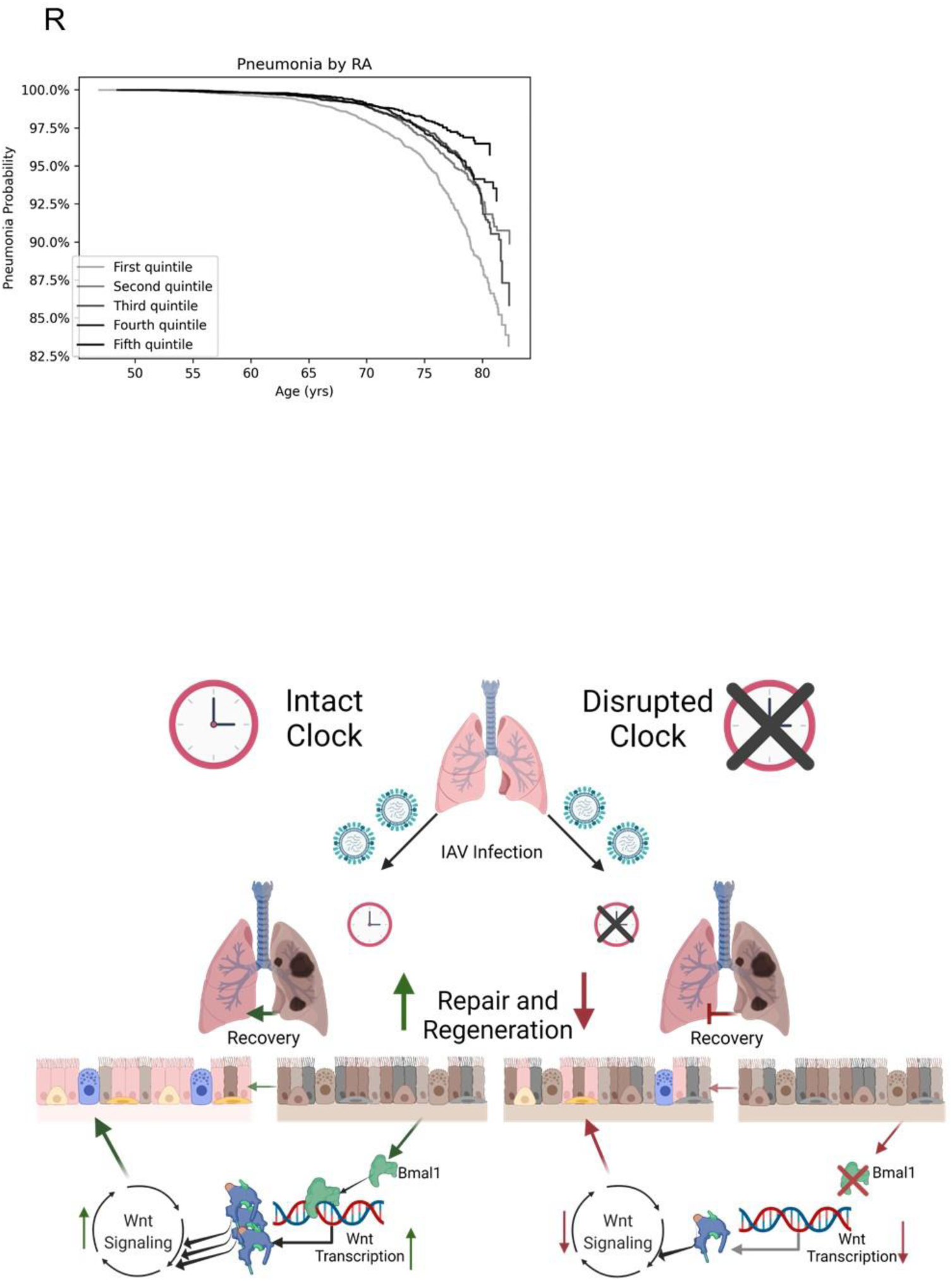

Therefore, we further investigated whether Wnt3a is a direct transcriptional target of *Bmal1* using chromatin immunoprecipitation (ChIP). Using bioinformatic analyses, we identified that the promoter region of Wnt3a contained three potential E-box motifs (CACGTG) to which BMAL1 was recruited. We then compared the fold enrichment between Wnt3a and Per2, a known Bmal1 target. The known *Bmal1* target gene *Per2* showed ∼ 12.54-fold enrichment of *Bmal1* recruitment to its promoter over IgG control (Fig 4C). Wnt3a was 6.12-fold enriched over its control, confirming that Bmal1 protein binds to the E-box motif in Wnt3a promoter (Fig 4C).

### Wnt3a treatment rescues *Bmal1^-/-^* phenotype in tracheal organoids

Next, we investigated whether activating Wnt signaling in the tracheal cells with disrupted clock would be sufficient to rescue the regenerative deficiency noted earlier. Treatment of the *Bmal1^-/-^* tracheal cells with Wnt3a resulted in a 2.2-fold increase in its regenerative capacity making it comparable to organoids grown from *Bmal1* sufficient cells (Fig 4D-G, *p<0.01, Kruskal-Wallis test). Similar results were noted in the *Bmal1*^creERT2/+^model wherein the regenerative deficiency was rescued with Wnt3a supplementation (Fig 5H-K, *p<0.01, ****p<0.0001, Kruskal-Wallis test). This was in contrast to alveolar organoids, wherein Wnt3a or ChIR supplementation did not rescue the regenerative loss associated with Bmal1 deficiency (S Fig 6I-Q., suggesting the role of additional circadian controlled mechanisms in AT2 regeneration.

Since tracheal organoids contain club cells, we used a club cell specific *Bmal1^-/-^* model to test if the disruption of the circadian clock in these cells results in worse repair following IAV infection. , We infected *Scgb1a1^Cre/+^: Bmal1^fl/fl^*mice at either CT11 or CT23 with IAV. We found that Scgb1a1Cre*^/+^: Bmal1^fl/fl^*had worse lung histology than *Scgb1a1^Creneg^:Bmal1^fl/fl^*mice. Interestingly, while the *Scgb1a1^Creneg^:Bmal1^fl/fl^*still maintained the time of day protection with the group infected at CT23 having significantly better histology than those infected at CT11, mice infected at subjective dusk or CT11 had significantly more alveolar and lung injury than mice infected at CT23 or subjective dawn (Fig 5L-M, P-Q, 20% vs 10% lung injury in zone 3: p<0.005, Mann-Whitney test; 4% vs 0.6% alveolar destruction in zone 4: p<0.01, Mann-Whitney test). This recapitulates the data seen with WT animals in Fig1B-D. However, among *Scgb1a1^Cre/+^: Bmal1^fl/fl^* this temporal protection was abolished and both groups had significantly more damage than the Cre^neg^ animals infected at CT23 (Fig 5N-O). Thus, loss of Bmal1 deficiency in club cells impairs lung repair and

### Individuals with poor circadian rhythms are more likely to have worse outcomes from pneumonia

To understand if this mechanistic work has relevance for human health, we extracted data from the UK Biobank. We have previously shown that poor circadian rhythmicity (defined as low rhythm amplitude or RA) was associated with increased risk of hospital admission for influenza and other lower respiratory tract infections^33^. RA was calculated based on hand worn actigraphy data and has been previously validated for the study of circadian rhythms^33^. These included hospital admission data that either followed or preceded the collection of the actigraphy data for defining circadian rhythms, which limits any causal inference. Therefore, here, our specific hypothesis was that poor circadian rhythmicity precedes the need for hospital admission for lung infection and thus can be considered a risk factor for the same. We use need for hospitalization as a surrogate marker for poor lung repair and regeneration here. Of the 64,789 participants who had actigraphy data recorded prior to a diagnosis of pneumonia, 1347 cases of pneumonia requiring hospital admission were listed. For the ease of comparison, we divided individuals into quintiles based on their RA. Among individuals with no prior history of pneumonia, low rhythmicity was significantly associated with high risk of future pneumonia diagnoses (Fig 5R with a -0.23 logHR per standard deviation RA, p = 2.8x10^-28^), while controlling for sex, ethnicity, overall health, income, history of smoking, age, BMI and college education. Kaplan-Meier curves for pneumonia separated by quintile of RA score (Fig 5R), not controlling for any cofactors reflect an increased risk in those with the poorest circadian rhythms. This finding provides important population level validation linking poor circadian rhythms with sub-optimal ability to repair and recover following lung infections.

## Discussion

In the current study, we used study the effect of circadian regulation on lung repair and regeneration, a process of incredible importance as we emerge from the current COVID-19 pandemic.

To the best of our knowledge, this is the first study linking the circadian clock with lung repair and regeneration. As seen for acute outcomes by our group^4, 9^ and others^2^, the circadian control of lung repair and regeneration manifested in a time-of-day specific protection. Mice infected at dawn had better recovery from IAV than those infected at dusk. Interestingly, the directionality of this protection is the opposite of what is observed vis-à-vis wound healing in the skin; wherein healing was better at night-time^5^.This finding suggests that circadian regulation is unique to each organ and underscores the importance of studying this specifically in the lung.

We demonstrated that epithelial cells from different levels of the respiratory tract with facultative stem cell function have a functional intrinsic clock and disruption of this circadian clock leads to loss of their regenerative potential. Further, the maturity of the lung organoids atl east from the tracheal paralleled the consolidation of circadian rhythms, suggesting that lung regeneration and clock function are intimately linked. We compared the lung transcriptome at the single cell level of both embryonic and postnatal *Bmal1* knockout models to dissect the novel signatures associated with loss of the circadian clock. Although there were many indications from the differential gene expression profile towards cell proliferation and differentiation, overall the result was subtle. However, this differential expression is likely an underestimation of the true functional effect of the clock following injury since the epithelium in an uninjured lung is rather quiescent and not actively proliferating. However, the use of uninfected lungs allowed us to circumvent the signals that emanate from the clock-gated regulation of the inflammation induced by IAV.

We focused on Wnt/β-catenin signaling pathway because of many reasons. First and foremost, activated canonical Wnt signaling has been shown to be crucial for proliferation, survival, and differentiation of lung epithelial stem/ progenitor cells^10, 20, 31, 34^. Further, the connection of the Wnt-pathways with the circadian clock in the maintenance of intestinal stem cell division has been shown to involve cyclical Wnt3A secretion from Paneth cells^6^. Similarly, Bmal1 specifically was found to act via the canonical Wnt signaling pathways to modulate adipocyte development and differentiation^32^. Finally, many of the downstream targets of the canonical Wnt pathway seem to oscillate across the 24 hr period, suggesting an upstream circadian influence on their regulation (Table 3). Here we found that upon disruption of the clock with *Bmal1* deletion, β-catenin was downregulated. Further, in tracheal organoids, the deficiency of regenerative capacity seen with *Bmal1* deletion could be completely rescued by exogenous administration of Wnt3a. Overall, these data confirm that circadian control over Wnt-driven lung regeneration. The data for AT2 organoids is a bit more complex. While there was loss of regenerative capacity in AT2 organoids from different models of clock disruption, the phenotype suggested a role for both the epithelium and mesenchyme and could not be rescued merely by exogenous administration of Wnt3a. However, the *in vivo* data clearly showed a loss of proliferative response in the absence of an intact AT2 clock as would be consistent with Wnt deficiency. Further studies will be needed to elucidate the exact mechanism underlying the clock control of lung repair and regeneration via AT2 cells, specifically how clock mechanisms in one epithelial compartment may affect other epithelial and mesenchymal compartments. However, despite this observation, across models of clock disruption, there is a consistent and unambiguous phenotype of deficient lung proliferation resulting in poor repair and regeneration following IAV.

The state of circadian health of an individual may be an important predictor of how one recovers following lung injury. This may have several implications for patients suffering from lung injury of various etiologies, well beyond IAV. Those with poorer rhythms are likely to have more protracted courses after infection. We have previously shown using the UK biobank database, that adults with less robust circadian rhythms were more likely to be hospitalized with influenza and other lower respiratory tract infections^33^. In the current study, we elaborated further on those analyses by showing that poor circadian rhythmicity preceded the diagnosis of pneumonia requiring hospital admission. These data attest to the importance of the circadian health of an individual as an important predictor of lung repair post lung injury. Most importantly, it provides powerful population-level evidence supporting the translational relevance of our mechanistic work.

In summary, through our current work, we introduce the circadian dimension to the incredibly complex process of lung repair and regeneration. Consideration of circadian context of lung injury and repair may not only provide novel therapeutic strategies but also allow for improved efficacy of many of our existing therapies towards enhancing lung recovery.

## METHODS

### Ethics statement

All animal studies were performed under guidance of the University of Pennsylvania Institutional Animal Care and Use Committee in accordance with institutional and regulatory guidelines.

### Mice

Embryonic *Bmal1* knockout mice (*Bmal1*^-/-^) where deletion of *Bmal1* prenatally disrupts clock-dependent oscillatory gene expression and behavioral rhythmicity and their littermate controls (*Bmal1*^+/+^) were generated by in-house breeding. AT2 cell-specific knockout of *Bmal1* (Sftpc^Cre-ERT2/+^:*Bmal1*^fl/fl^) was generated by crossing Sftpc^Cre-ERT2/+^, a tamoxifen-inducible Cre, with *Bmal1*^fl/fl^mice^9^ .To generate postnatal *Bmal1* knockout mice, 2 months old (unless specified) *Bmal1*^creERT2/+^mice were treated with 5mg (in 50µl) of tamoxifen via oral gavage, for 5 consecutive days. Tamoxifen was reconstituted to 100mg/ml solution with ethanol and corn oil and thawed at 55°C prior to administration. *Bmal1*^creERT2neg^, Sftpc^CreERT2neg^: *Bmal1*^fl/fl^littermates treated with tamoxifen served as controls^4^. Club cell specific *Bmal1* knockout (Scgb1a1^Cre/+^: *Bmal1*^fl/fl^) was generated by crossing CCSP-cre, with *Bmal1*^fl/fl^ mice, Scgb1a1^Creneg^:*Bmal1*^fl/fl^served as controls^4^. PER2::LUC mice (B6.129S6-Per2^tm1Jt^/J), in which the luciferase gene is fused in-frame to the 3′ end of the endogenous mPer2 gene (one of the core clock genes), were purchased from Jackson Laboratory animal facility (stock: 006852). *Cry1^-/-^Cry2^-/-^(Cry1,2 DKO)* mice were a gift from Dr. Amita Sehgal^35, 36^. Male and female mice were used in approximately equal proportion for all experiments.

### Lung Injury

For influenza infections, mice were lightly anesthetized with isoflurane and infected intranasally (i.n.) with sub-lethal dose of IAV (H1N1:PR8, dose: 10 PFU), in a volume of 40 µl. Mice were infected at either ZT11 (an hour before onset of activity phase or dusk) or ZT23 (an hour before onset of rest phase or dawn). By circadian nomenclature ZT0 refers to time at which lights went on. To remove any confounding effect of light-driven behavioral rhythms not entirely reflective of the endogenous circadian clock, all transgenic mice were infected under constant darkness conditions. The times corresponding to ZT11 and ZT23 in constant darkness is referred to as CT11 (subjective dusk) or CT23 (subjective dawn) respectively, by circadian convention. Weight and clinical scores were recorded at serial intervals as described previously^4, 9^. Lungs were harvested on either day 8 or day 30, based on the experimental question.

### Single cell preparation, and sorting

Tracheal organoids: Mice tracheas were processed by peeling off the epithelium and digesting in a 1:1 mixture of DNAse II (Roche) and Liberase (Roche) for 30 min at 37°C. After centrifugation at 1200 rpm, the cell pellet was re-suspended in 0.25% trypsin EDTA and incubated at 37 °C for 10 min. Thereafter the cell suspension was filtered through a 40 µm cell strainer (BD Biosciences) and recovered cell pellet was plated in the appropriate media.

Bronchial epithelial organoids: Lungs were harvested after PBS perfusion followed by digestion with DNAse II and Liberase at 37°C for 20 mins. Dissociated lung tissue was passed through a 70 µm cell strainer (BD Biosciences), followed by centrifugation and RBC lysis. After magnetic bead-based depletion of leukocytes and endothelial cells (CD45^-^CD31^-^), the single cell suspension was sorted for alveolar epithelial cells (AT2 as DARQ7^-^EPCAM^+^CD104^-^) and CD104^+^bronchial epithelial cells (DARQ7^-^EPCAM^+^CD104^+^) on BD FACSJazz™ (S. Fig 1A). Using this strategy, the alveolar fraction, is likely to contain some AT1 along with AT2 cells and the CD104+ fraction has a mix of bronchial epithelial cells. CD104^-^ Epcam+ population was used as a surrogate for AT2 organoids. This was confirmed on histology. Similarly, the CD104^+^organoids, represented the regenerative capacity of a mix of basal and club cells.

Lung fibroblasts for organoid assays were isolated from embryonic *Bmal1*^-/-^, wild type littermate *Bmal1*^+/+^ controls, *Bmal1*^creERT2/+^, and *Bmal1*^creERT2neg^littermate controls. To do so, lungs were harvested, digested, and processed into a single cell suspension as described above. Thereafter they were plated and serially passaged in DMEM/F12 (11320033 Life Technologies) supplemented with 10% FBS and penicillin-streptomycin (P433310,000 U/mL, Sigma) 3 times.

### Single Cell RNA sequencing

Lungs from embryonic *Bmal1*^-/-^, *Bmal1*^+/+^ controls, *Bmal1*^creERT2/+^, and *Bmal1*^creERT2neg^littermates were dissected, and single-cell preparation was obtained as described above. Single cell suspension was sorted as DARQ7^-^CD31^-^CD45^-^ lung cells (S. Fig 1B). The sorted cells were loaded onto a Chromium Controller instrument (10× Genomics, Pleasanton, CA, USA) to generate single-cell barcoded droplets (GEMs) according to the manufacture’s protocol using the 10× Single Cell 3’ v1 chemistry. The resulting libraries were uniquely indexed using the Chromium i7 Sample Index Kit, pooled, and sequenced were sequenced on the Illumina NovaSeq 6000 sequencer using an SP 100 cycles flow cell in a paired-end, single indexing run. Sequencing for each library targeted 25,000 mean reads per cell. Data was then processed using the Cell ranger pipeline (10x genomics, v.3.1.0) for demultiplexing and alignment of sequencing reads to the mouse mm10 transcriptome and creation of feature-barcode matrices. Individual single cell RNAseq libraries were aggregated using the cell ranger aggr pipeline. Libraries were normalized for sequencing depth across all libraries during aggregation.

### Chromatin Immunoprecipitation

Lungs were cut into small pieces and cross-linked with 1% formaldehyde for 10 min at room temperature (RT). 125mM of glycine (Thermo Fisher Scientific) was added to stop the crosslinking reaction with additional incubation for 5 minutes on the revolver at RT. Tissue was homogenized for 10s at 06m/sec on MP Fastprep-24 5G (MP Biomedicals). Cell pellet was washed with cold PBS, centrifuged, and then suspended in swelling buffer (5 mM PIPES pH 8.0, 85 mM KCl, 1% NP40, and protease inhibitor cocktail) for 30 minutes on ice. The crude nuclear preparation was centrifuged, and nuclei were suspended in nuclear lysis buffer (50 mM Tris-HCl pH 8.0, 10 mM EDTA, 1% SDS, and protease inhibitor cocktail) and sonicated to an average length of about 300-500 base pairs using Diagenode Bioruptor (UCD-200), at 4°C on setting of “3” for 35 minutes.

Sonicated samples were diluted 10-fold with IP dilution buffer (16.7 mM Tris-HCl pH 8.0, 0.01% SDS, 1.1% Triton X-100, 1.2 mM EDTA, 167 mM NaCl, and protease inhibitor cocktail) and incubated with anti-Bmal1 antibody (5 µg, ab3350, abcam) overnight at 4°C. DNA complexes were collected on Dynabeads protein A® (1001D, Thermofisher) and serially washed once with dialysis buffer (50 mM Tris-HCl pH 8.0, 2 mM EDTA, and 0.2% sarkosyl), three times with IP wash buffer (100 mM Tris-HCl pH 9.0, 500 mM LiCl, 1% NP40, 1% Deoxycholate, and protease inhibitor cocktail), and finally in TE buffer (1M TrisCL, 05.M EDTA, pH 8.0). Samples were eluted with elution buffer (50 mM NaHCO_3_ and 1% SDS). Crosslinking was reversed by overnight incubation with 0.3 M NaCl at 65°C followed by proteinase K treatment (0.5 M EDTA, 1 M Tris-HCl (pH 7.5), and 10 mg/mL proteinase K) and DNA clean up (MiniElute reaction clean up kit, 28204, Qiagen). qPCR was performed using mPer2 and Wnt3a primers. Rabbit IgG antibody (CS200581, Millipore) was used as a negative control and anti-Histone H3 (tri methyl K9, AB6002, Abcam) antibody was used as a positive control. Data was analyzed using % Input method.

### Western Blot Analysis

Total protein from lungs were extracted. 20–50 µg of protein from each sample was resolved on SDS-PAGE gels followed by immunoblotting after PVDF membrane transfer, probed with β-catenin (1:500, 610154 BD Biosciences), β actin (1:3000, ab8227 abcam), and secondary antibodies anti-rabbit HRP-linked IgG (1:5000 7074P2, Cell signaling) or anti-mouse IgG (1:5000 A2228 Sigma) and developed by chemiluminescence (34579 Supersignal; Thermo Scientific). Relative changes in the expression levels of interested proteins were measured by densitometry analysis using ImageJ 1.47 software and normalized to β-actin.

### Ex vivo Organoid assays

Tracheal cells (60,000-100,000 cells/ well in 96 well plate) were cultured in small airway growth media (SABM^TM^Basal Medium, CC-3119 Lonza) containing 1× insulin, 1x transferrin,1x hEGF, and 1x bovine pituitary extract (SAGM^TM^ SingleQuots^TM^ supplements, CC-4124 Lonza), 0.1 µg/ml Cholera Toxin (C9963 Sigma), 0.01µM Retinoic acid (Y0503 Sigma) and 5% FBS (Denville) on growth factor–reduced, phenol-free matrigel (356231 Corning).

Equal number of CD104^+^basal cells sorted from distal lung, were cultured in C12 media^40^ supplanted with growth factors onto growth factor–reduced, phenol-free matrigel depending on the yield of cells per experiment per experimental condition in 96 well plate.

For alveolar organoid assays: For each experiment, 5 × 10^3^Epcam^+^CD104^-^were isolated as described above and mixed with 5 x10^4^ lung fibroblasts. Cells were then suspended media similar to tracheal organoids and growth factor-reduced, phenol-free matrigel at 1:1 concertation. 90μl of the cell/media/matrigel mixture was then aliquoted into individual 24-well cell culture inserts and allowed to solidify at 37^0^C. Complete SAGM was added to each well. Rock inhibitor Y27632 (Sigma) was included in the media for the first two days. Cultures were maintained at 37^0^C, 5% CO2. Media was replenished every 48h until day 8 for tracheal and CD104+organoids, and day 21 for AT2 organoids.

Ligand treatments of tracheal and AT2 organoids were performed using the following reagents at the indicated concentrations: Wnt3a 200ng/ml (315-20 Peprotech), Fgf10 50ng/ml (100-26 Peprotech), CHIR99021 3 µm (72052 StemCell Technologies), DMSO (final concentration 0.05%, D2650 Sigma) was used a control.

Each organoid assay was run with at least three technical replicates. Images were recorded using EVOS M7000 imaging system (Thermo Fisher Scientific). Colony forming efficiency (CFE: number of colonies formed/number of cells plated/well) of organoids was determined using a customized macro in Image J 1.47.

### Real-time bioluminescence recording of PER2::LUC organoids

For luciferase assay, tracheal cells, CD104^+^ distal lung cells or AT2s were harvested from *mPer2^luc^* mice (Stock No: 006852). In half of the wells, the bioluminescence was recorded without any synchronization agents. For other wells, starting seven days post seeding, tracheal organoids were then pulsed with either 10µM dexamethasone (Sigma) in SAGM, and CD104^+^ organoids in C12 media for synchronization. Similarly, on day 20 post seeding, AT2 organoids were pulsed with dexamethasone in SAGM. Bioluminescence outputs were recorded by adding beetle luciferin potassium salt (E1602 Promega) to organoid media at a final concentration of 10 mM. Cultures were monitored for light output using a custom-made bioluminescence recording system (Cairn Research Ltd, UK) composed of charge-coupled device camera (Andor iKon-M 934) mounted on the top of an Eppendorf Galaxy 170R CO2 incubator^37^.

### Histology

Lungs harvested at the end of the experiment were fixed by inflation with 10% buffered formalin at 20 mm H_2_O of pressure, paraffin embedded, and stained with H&E stain. Stained slides were digitally scanned at 40x magnification using an Aperio CS-O slide scanner (Leica Biosystems, Chicago IL). Representative images were taken from scanned slides using Aperio ImageScope v12.2.2.5015 (Leica Biosystems, Chicago, IL). Three-four random fields per lung section were imaged for Ki67 scoring. The histological and cytological scoring were performed in a blinded fashion. Numerical codes were used to identify these slides during the scoring. Once all the data were recorded, the identity was unmasked, and final analyses undertaken according to the study group.

Similarly, after 8 days for tracheal and CD104^+^, and 21 days for AT2 organoids were fixed in 2% paraformaldehyde, embedded in Histogel (Richard-Allen), dehydrated, paraffin embedded, and sectioned. Hematoxylin and eosin staining was performed to examine the whole lung and organoid morphology.

Immunofluorescence was performed following heat antigen retrieval methods and stained with the following antibodies. Sftpc (1:100, anti Pro Surfactant protein, rabbit, AB3786 Millipore), Ki67 (1:200, mouse, 550609 BD Biosciences), Podoplanin (1: 100 University of Iowa Developmental Studies Hybridoma Bank clone 8.1.1). Podoplanin signal intensity quantification using a three random region of interests were analyzed using LasX software (Leica Microsystems) for images taken over the course of the experiment.

### UK biobank Analyses

UK biobank is a large-scale database which contains biomedical information from >500,000 participants from UK who are followed longitudinally. The data is available for researcher across the world. (https://www.ukbiobank.ac.uk). For the purpose of this analyses we defined pneumonia as the following ICD10 codes which required admission to the hospital: (1) Pneumonia: B58.3 J10.0 J11.0 J16.8 J17.2 J18 J18.8 J18.9; (2) Bacterial pneumonia: A20.2 A21.2 A22.1 A31.0 A42.0 A43.0 A48.1 A78 B96.0 B96.1 J14 J15.0 J15.2 J15.3 J15.4 J15.5 J15.6 J15.7 J15.8 J15.9 J16.0; (3) Pneumococcal pneumonia: J13 J18.1; (4) Pseudomonal pneumonia: J15.1; (5) MRSA pneumonia: none; (6) Viral pneumonia: B01.2 B05.2 B25.0 J12 J12.0 J12.1 J12.2 J12.8 J12.; (7) Pneumonia due to fungus (mycoses): (8) B20.6 B37.1 B38.0 B38.1 B38.2 B44.0 B59 J17; (9) Bronchopneumonia and lung abscess: J18.0 J85 J85.0 J85.1 J85.2 J85.3. The robustness of circadian rhythmicity in participants was based on their actimetry data obtained from hand worn devices. In particular Relative Amplitude (RA) was defined as relative difference in day versus night: (day – night) / (day + night). Acceleration-derived RA (acceleration) goes from 0 to 1 with 1 being “the best”, perfect day/night differences. The RA of the rhythms was divided into quintiles and Cox Proportional Hazards model was used to test the effect of circadian rhythmicity on the likelihood of later to be admitted to the hospital with a diagnosis of pneumonia.

### Statistics

All statistical analyses were performed using GraphPad (Prism V8). For experiments with more than 2 groups and normally distributed data, a one-way ANOVA was performed, and p value adjustments for multiple comparisons were performed using Dunnett HSD. In experiments where the data was not normally distributed, Mann-Whitney, and Kruskal-Wallis tests were performed. Bioluminescence data traces were analyzed using a modified version of the R script “CellulaRhythm”. Statistical data was considered significant if P < 0.05. All the plots represent mean of biological replicates, and error bars represent standard error of mean.

### Statement on rigor and reproducibility

All studies were performed using animals from either Jackson Labs and animals from in-house breeding. The background strain of each genetically modified animal has been specified and controls were cre negative littermates on that same background. Animals for ScSeq and all the organoid assays were sacrificed between ZT1-ZT2 unless otherwise stated. Reported findings are summarized results from 3-6 independent experiments.

## Acknowledgements

We are thankful to members of the FitzGerald lab-- Dr. S. Teegarden and Taylor Hollingsworth for help with animal breeding and general lab management. Dr. A.S. Sase for help with chromatin immune precipitation assay. Dr. P. Chandrasekaran for immunofluorescence techniques. Samuel Gavronski and Aastha Bansal undergraduate student interns in Sengupta lab for help with image analyses. This work was supported by the NHLBI-K08HL132053 (SS), University of Pennsylvania (SS)

## Author Contributions

SS conceived the project; AN, SS, EEM, GSW and AS designed experiments; AN, KF, YI, UV, DBF and SS performed experiments and collected data. AN, AB, MM, NL and TB analyzed data. ABR and DBF were involved in the interpretation of data. GAF, GG and TB were involved in the UK biobank analyses; AS also provided *Cry1,2 DKO* mice. AN wrote original draft with help from SS. SS supervised all research activities.

## Data Availability

The RNA-Sc-seq data is available at the NCBI GEO accession number???. All other data is available as source data.

## Competing interest statement

The authors declare no competing interests.

---

**Figure S1.**
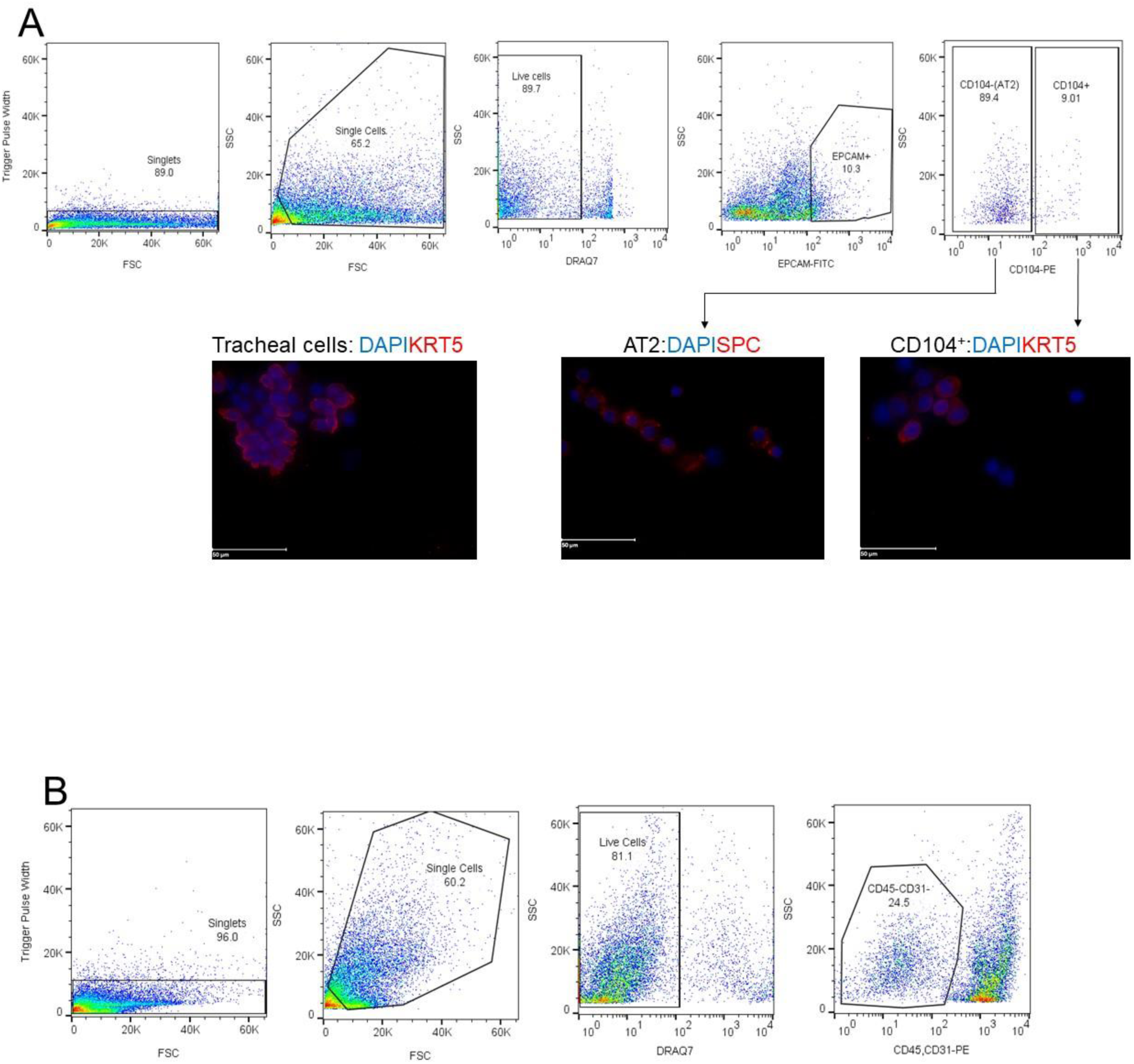

**Figure S2.**
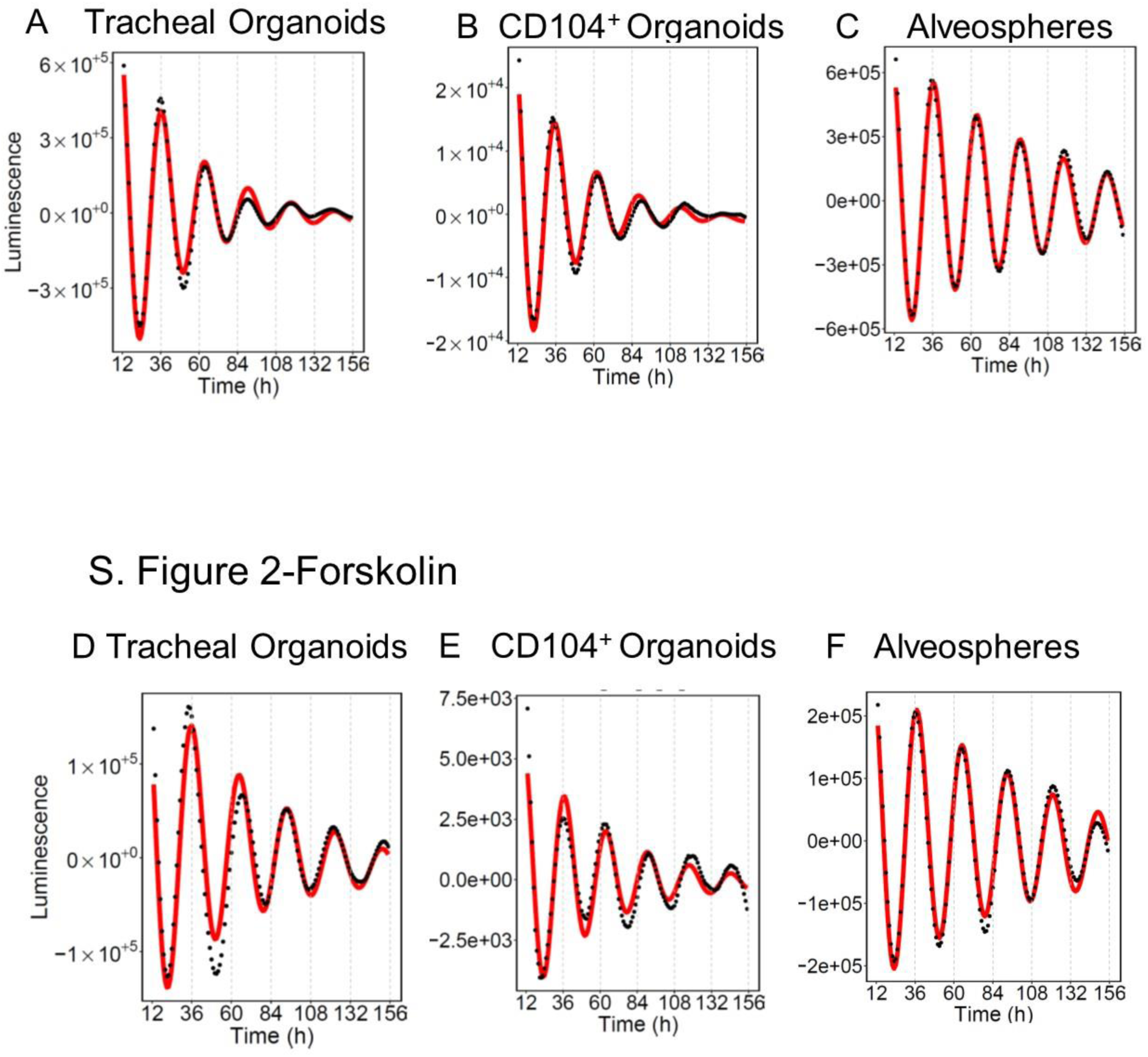

**Figure S3.**
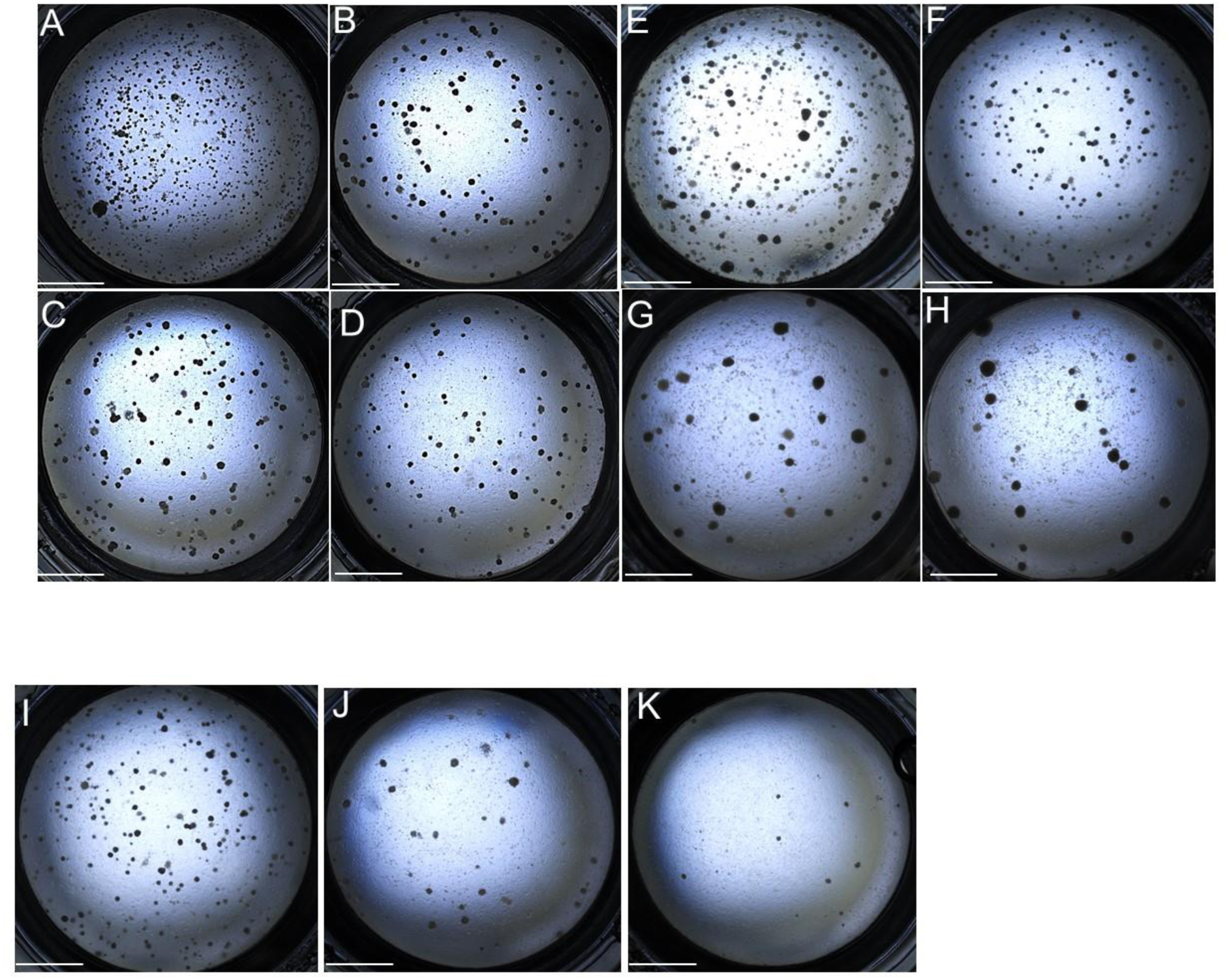
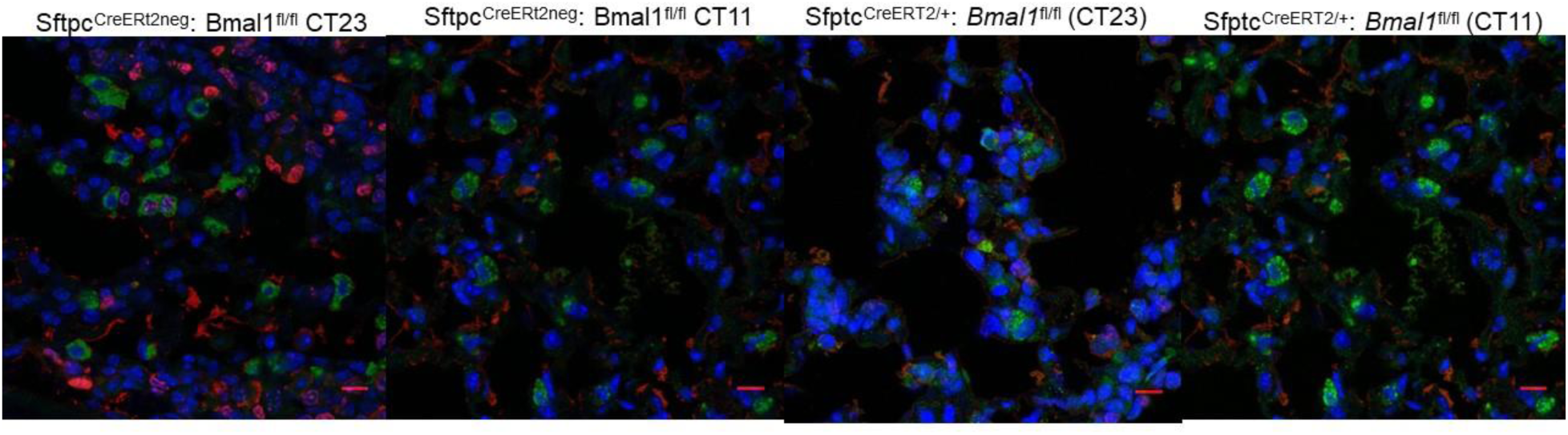

**Figure S4.**
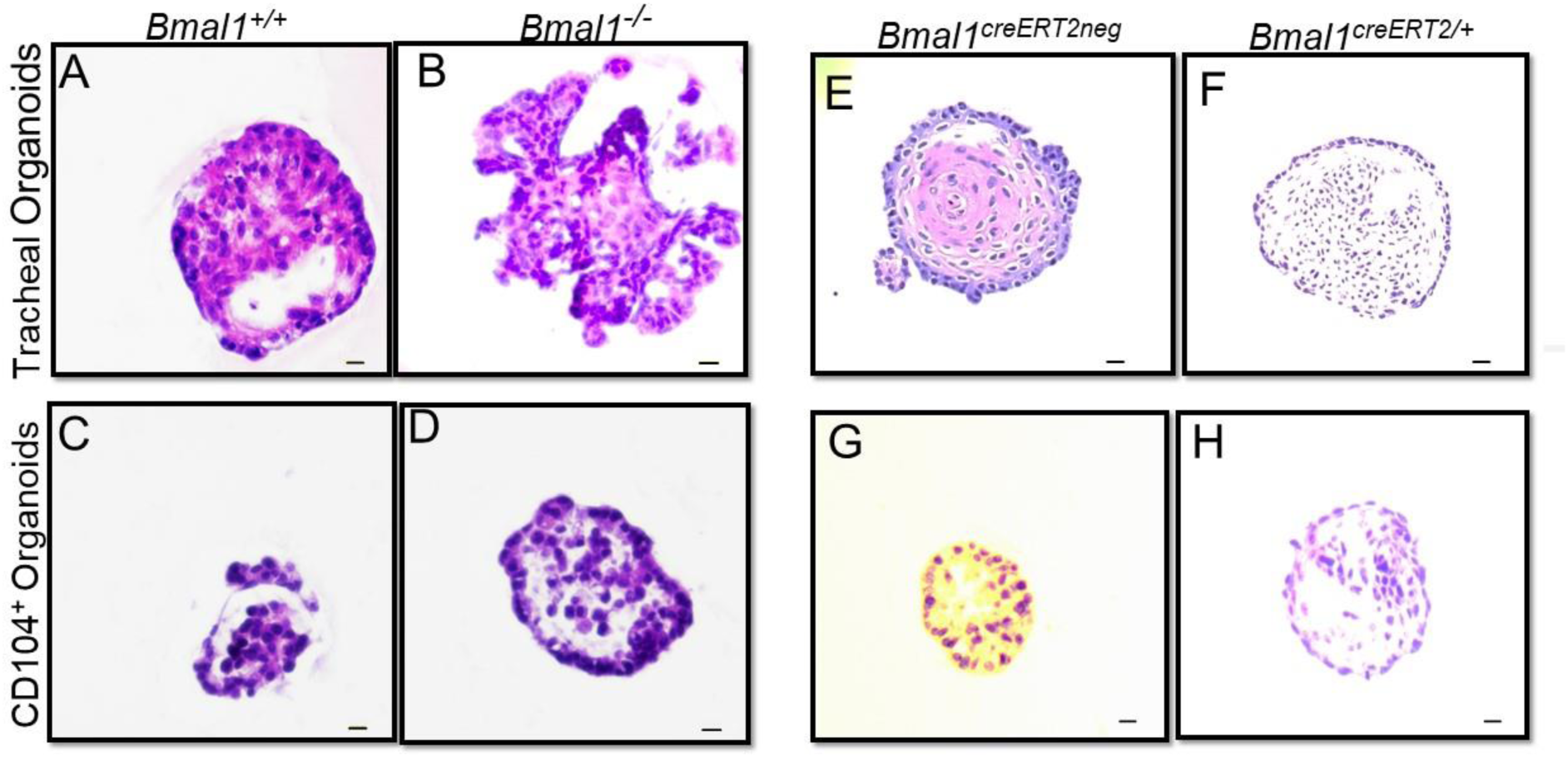

**Figure S5.**
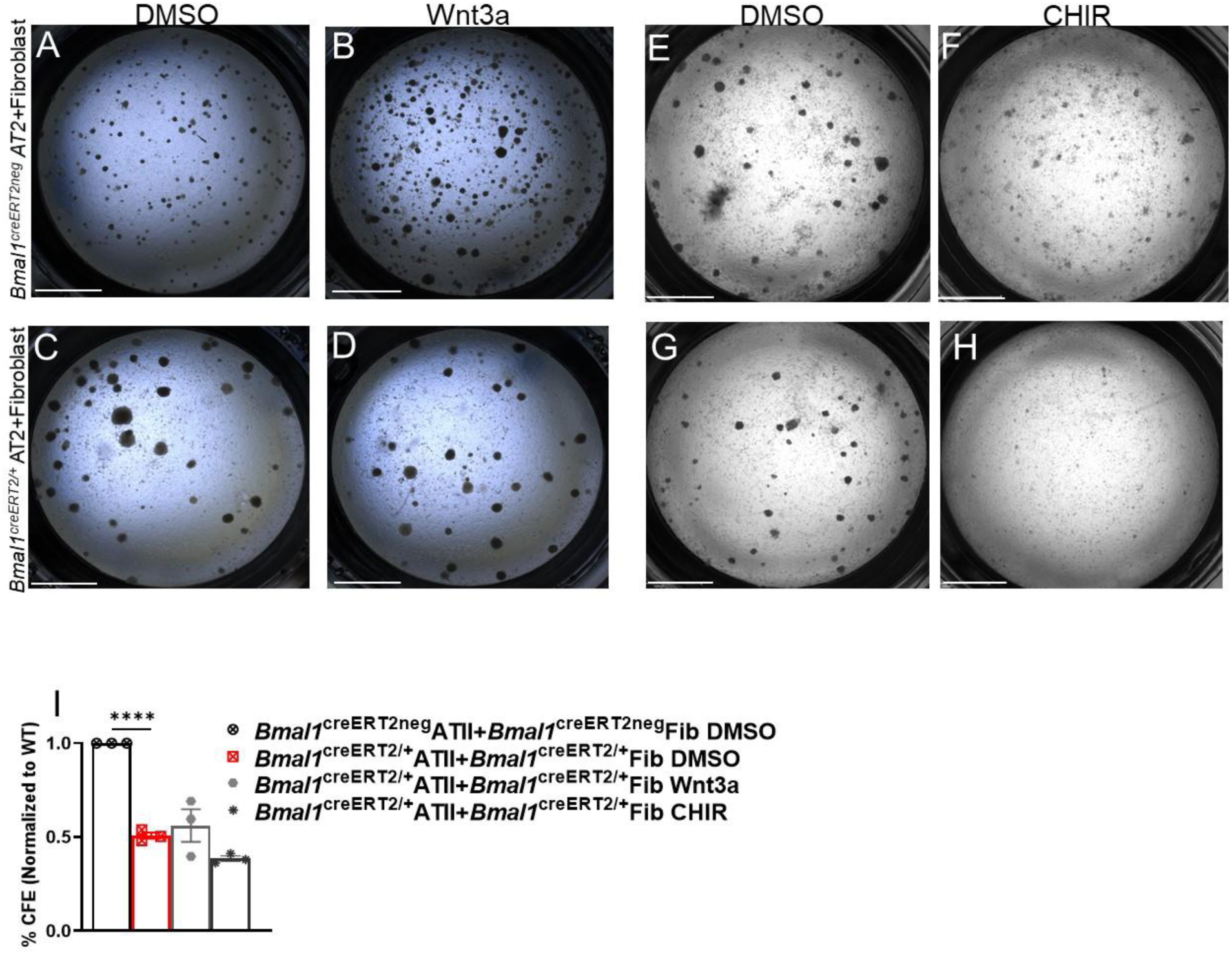

## Notes

### Competing Interest Statement

The authors have declared no competing interest.

## REFERENCES

1. Takahashi, J.S. Transcriptional architecture of the mammalian circadian clock. Nat Rev Genet 18, 164–179 (2017).

2. Ehlers, A., et al. BMAL1 links the circadian clock to viral airway pathology and asthma phenotypes. Mucosal Immunol 11, 97–111 (2018).

3. Sukumaran, S., Jusko, W.J., Dubois, D.C. & Almon, R.R. Light-dark oscillations in the lung transcriptome: implications for lung homeostasis, repair, metabolism, disease, and drug action. J Appl Physiol (1985) 110, 1732–1747 (2011).

4. Sengupta, S., et al. Circadian control of lung inflammation in influenza infection. Nat Commun 10, 4107 (2019).

5. Hoyle, N.P., et al. Circadian actin dynamics drive rhythmic fibroblast mobilization during wound healing. Sci Transl Med 9(2017).

6. Matsu-Ura, T., et al. Intercellular Coupling of the Cell Cycle and Circadian Clock in Adult Stem Cell Culture. Mol Cell 64, 900–912 (2016).

7. Stokes, K., et al. The Circadian Clock Gene BMAL1 Coordinates Intestinal Regeneration. Cell Mol Gastroenterol Hepatol 4, 95–114 (2017).

8. Major, J., et al. Type I and III interferons disrupt lung epithelial repair during recovery from viral infection. Science 369, 712–717 (2020).

9. Issah, Y., et al. Loss of circadian protection against influenza infection in adult mice exposed to hyperoxia as neonates. Elife 10(2021).

10. Zacharias, W.J., et al. Regeneration of the lung alveolus by an evolutionarily conserved epithelial progenitor. Nature 555, 251–255 (2018).

11. Alvarez-Dominguez, J.R., et al. Circadian Entrainment Triggers Maturation of Human In Vitro Islets. Cell Stem Cell 26, 108–122 e110 (2020).

12. Moore, S.R., et al. Robust circadian rhythms in organoid cultures from PERIOD2::LUCIFERASE mouse small intestine. Dis Model Mech 7, 1123–1130 (2014).

13. Barkauskas, C.E., et al. Type 2 alveolar cells are stem cells in adult lung. J Clin Invest 123, 3025–3036 (2013).

14. Chapman, H.A., et al. Integrin alpha6beta4 identifies an adult distal lung epithelial population with regenerative potential in mice. J Clin Invest 121, 2855–2862 (2011).

15. Hogan, B.L., et al. Repair and regeneration of the respiratory system: complexity, plasticity, and mechanisms of lung stem cell function. Cell Stem Cell 15, 123–138 (2014).

16. Zuo, W., et al. p63(+)Krt5(+) distal airway stem cells are essential for lung regeneration. Nature 517, 616–620 (2015).

17. Vaughan, A.E., et al. Lineage-negative progenitors mobilize to regenerate lung epithelium after major injury. Nature 517, 621–625 (2015).

18. Ray, S., et al. Rare SOX2(+) Airway Progenitor Cells Generate KRT5(+) Cells that Repopulate Damaged Alveolar Parenchyma following Influenza Virus Infection. Stem Cell Reports 7, 817–825 (2016).

19. Yang, G., et al. Timing of expression of the core clock gene Bmal1 influences its effects on aging and survival. Sci Transl Med 8, 324ra316 (2016).

20. Frank, D.B., et al. Emergence of a Wave of Wnt Signaling that Regulates Lung Alveologenesis by Controlling Epithelial Self-Renewal and Differentiation. Cell Rep 17, 2312–2325 (2016).

21. Angelidis, I., et al. An atlas of the aging lung mapped by single cell transcriptomics and deep tissue proteomics. Nat Commun 10, 963 (2019).

22. Xie, T., et al. Single-Cell Deconvolution of Fibroblast Heterogeneity in Mouse Pulmonary Fibrosis. Cell Rep 22, 3625–3640 (2018).

23. Hecht, A., Vleminckx, K., Stemmler, M.P., van Roy, F. & Kemler, R. The p300/CBP acetyltransferases function as transcriptional coactivators of beta-catenin in vertebrates. EMBO J 19, 1839–1850 (2000).

24. Shtutman, M., et al. The cyclin D1 gene is a target of the beta-catenin/LEF-1 pathway. Proc Natl Acad Sci U S A 96, 5522–5527 (1999).

25. Tetsu, O. & McCormick, F. Beta-catenin regulates expression of cyclin D1 in colon carcinoma cells. Nature 398, 422–426 (1999).

26. Lai, K.K.Y., et al. Stearoyl-CoA Desaturase Promotes Liver Fibrosis and Tumor Development in Mice via a Wnt Positive-Signaling Loop by Stabilization of Low-Density Lipoprotein-Receptor-Related Proteins 5 and 6. Gastroenterology 152, 1477–1491 (2017).

27. Shimura, T., et al. Implication of galectin-3 in Wnt signaling. Cancer Res 65, 3535–3537 (2005).

28. Shimura, T., et al. Galectin-3, a novel binding partner of beta-catenin. Cancer Res 64, 6363–6367 (2004).

29. Song, S., et al. Galectin-3 mediates nuclear beta-catenin accumulation and Wnt signaling in human colon cancer cells by regulation of glycogen synthase kinase-3beta activity. Cancer Res 69, 1343–1349 (2009).

30. Giangreco, A., et al. beta-Catenin determines upper airway progenitor cell fate and preinvasive squamous lung cancer progression by modulating epithelial-mesenchymal transition. J Pathol 226, 575–587 (2012).

31. Nabhan, A.N., Brownfield, D.G., Harbury, P.B., Krasnow, M.A. & Desai, T.J. Single-cell Wnt signaling niches maintain stemness of alveolar type 2 cells. Science 359, 1118–1123 (2018).

32. Guo, B., et al. The clock gene, brain and muscle Arnt-like 1, regulates adipogenesis via Wnt signaling pathway. FASEB J 26, 3453–3463 (2012).

33. Sengupta, S., Brooks, T.G., Grant, G.R. & FitzGerald, G.A. Accounting for Time: Circadian Rhythms in the Time of COVID-19. J Biol Rhythms 36, 4–8 (2021).

34. Lee, J.H., et al. Anatomically and Functionally Distinct Lung Mesenchymal Populations Marked by Lgr5 and Lgr6. Cell 170, 1149–1163 e1112 (2017).

35. Thresher, R.J., et al. Role of mouse cryptochrome blue-light photoreceptor in circadian photoresponses. Science 282, 1490–1494 (1998).

36. Zhang, S.L., et al. A circadian clock regulates efflux by the blood-brain barrier in mice and human cells. Nat Commun 12, 617 (2021).

37. Rey, G., et al. The Pentose Phosphate Pathway Regulates the Circadian Clock. Cell Metab 24, 462–473 (2016).

