## Supplementary material for "Circadian regulation of lung repair and regeneration": Figure legends

**Figure 1: Time of day infection affects long term lung repair and regeneration**

A) Schematic of experimental design for long term recovery from Influenza Virus A (IAV) infection Two groups C57Bl6 mice aged 8-10 weeks were maintained in 12 h light: dark (LD) cycles. They were infected with intranasal IAV (sub lethal dose of H1N1 (PR8); intransal-i.n.) at either the start of the light cycle (ZT23; ZT0 being the time at which light go on in a 12 h LD cycle) or at the start of the dark cycle (ZT11), and subsequently recovered to day 30.

B) Influenza-induced lung injury was categorized into the following groups: Zone 1-minimal injury, Zone 2-minor injury with mild interstitial thickening, Zone 3- severe alveolar injury or Zone 4-complete alveolar destruction. Representative micrographs of H&E-stained lung sections on day 30 post-infection quantified as defined above.

C) Severity of lung injury quantified using the objective histopathological scoring system defined above by a researcher blinded to study group (n= 9-10 mice per circadian time point, ** $p<0.007$, *$p<0.02$Mann-Whitney test, each data point represents individual animal, data pooled from two independent experiments)

D) Schematic of experimental design for organotypic assay. After magnetic bead-based depletion of leukocytes and endothelial cells ($CD45^-CD31^-$), the single cell suspension was sorted and plated to develop alveolar type 2 (AT2) organoids from as $DRAQ7^-EPCAM^+CD104^-$ or $CD104^+$ organoids from $DRAQ7^-EPCAM^+CD104^+$ cells. Trachea were also digested, and mixed tracheal cells plated for tracheal organoids.

Representative records of bioluminescence showing circadian profiles of PER2 expression in organoids grown from different levels of respiratory tree from *mPer2*$^{Luc}$ knockin mice. E) tracheal organoids (day 7 onwards in SAGM) F) $CD104^+$ organoids (day 7 onwards C12) G) AT2 organoids (day 18 onwards in SAGM). H) Bioluminescence recording of tracheal organoids 4 days post seeding. Data summarized from two independent experiments. Bioluminescence data traces were analyzed with a modified R script CellulaRhythm.

**Figure 2: Deletion of *Bmal1* results in reduced regenerative capacity in Lung organoids**

Lungs from embryonic *Bmal1*$^{-/-}$ knockouts and their Bmal1$^{+/+}$ littermates were used for organotypic assays. For postnatal knockouts, *Bmal1*$^{creERT2/+}$ and their Cre$^{neg}$ littermates were treated with tamoxifen at 8 weeks of age. Representative images of tracheal organoids from cells harvested from (A) *Bmal1*$^{+/+}$ B) *Bmal1*$^{-/-}$ C) *Bmal1*$^{creERT2neg}$ D) *Bmal1*$^{creERT2/+}$ mice; E-F) Colony forming efficiency (CFE) of each condition.

Representative images of CD104$^+$ distal lung cell organoids grown from cells harvested from G) *Bmal1*$^{+/+}$ wild type littermates H) *Bmal1*$^{-/-}$ I) *Bmal1*$^{creERT2neg}$ J) *Bmal1*$^{creERT2/+}$ K,L) CFE.

L-M) Representative images of tracheal organoids from *Cry1*$^{-/-}$*Cry2*$^{-/-}$ (*Cry1,2* DKO).

N-O) Quantification expressed as CFE of AT2 organoids grown from embryonic and tamoxifen treated postnatal *Bmal1* knockout mice, P-Q) AT2 organoids co-cultured with *Cry1,2* DKO fibroblasts

4-5 independent experiments with 3 technical replicates. scale bar: 2000μm. Compiled data are expressed as mean ± SEM. E; ***p<0.0005, F,K; *<0.01, L; *<0.04, Unpaired t test with Welch's correction. N; #p=0.06, Unpaired t test with Welch's correction, N; *p<0.05, O; *p<0.01, **p<0.005, Ordinary one-way ANOVA. P; **<p0.037 ****p<0.0001, Q; *p<0.01, ****p<0.0001 Ordinary one-way ANOVA.

**Figure 3: Single-cell transcriptome analysis of *Bmal1*$^{-/-}$ lung cells reveal downregulation of Wnt associated pathways**

Single-cell transcriptomic analysis was performed on integrated cells from *Bmal1*$^{+/+}$, *Bmal1*$^{-/-,}$ *Bmal1*$^{creERT2neg}$, and *Bmal1*$^{creERT2/+}$ mice. A) Uniform manifold approximation and projection (UMAP) visualization dimension reduction analysis of scRNA-seq data generated using a Seurat pipeline B) The dot plot showing the percentage of cells expressing the respective selected marker gene based on dot size C) Venn diagram depicting the number of differentially expressed genes D) GO biological process enrichment of differentially expressed genes in both the knockout models E) Violin plots showing expression levels of differentially expressed genes.

**Figure 4: Disruption of the circadian clock lead to diminished proliferation post IAV infection**

Representative images for A) Sftpc and Pdpn staining of mice from experimental design explained in Fig 1A B) with quantitation expressed as absolute Sftpc$^+$ AT2 cells and fluorescence intensity of Pdpn$^+$ AT1 cells. Scale Bar: 1000μm, 40x (n=9, unpaired t test with Welch’s correction, data pooled from two independent experiments).

*Sftpc*$^{CreERt2/+}$*: Bmal1*$^{fl/fl}$ mice (AT2 specific *Bmal1* KO) and their cre$^{neg}$ littermates were treated with tamoxifen at 6–8 weeks of age. Mice including *Cry* 1, 2 DKO and Scgb1a1$^{Cre/+}$: *Bmal1*$^{fl/fl}$ (club cell specific *Bmal1* KO) and their Cre$^{neg}$ littermates were acclimatized to reverse cycles of 12 hr LD for 2 weeks. Thereafter, they were moved to constant darkness (DD) 2 days prior to administering

IAV at either CT23 or CT11, maintained in DD until 8 days post infection. Representative Ki67 staining images from C) Wild type, D) *Cry* 1, 2 DKO; E-F) Scgb1a1$^{Creneg}$*:Bmal1*$^{fl/fl;}$ G-H) Scgb1a1$^{Cre/+}$: *Bmal1*$^{fl/fl}$ ; J-K) *Sftpc*$^{CreERT2neg}$*:* Bmal1$^{fl/fl}$ ;L-M) *Sftpc*$^{CreERt2/+}$*:Bmal1*$^{fl/fl}$ mice. E, I, N) quantitation expressed as percentage Ki67$^{+}$ cells/5 random fields.

Compiled data are expressed as mean ± SEM. Each data point represents individual animal, n=3-4 mice/ condition, scale bar: 100µm, E; *p<0.0129*,* unpaired t test, Welch's correction. I; **p<0.003, Ordinary one-way ANOVA. N; **p<0.004*,* p*<0.05, Ordinary one-way ANOVA.

**Figure 5: Activation of Wnt signaling in tracheal organoids rescues the regenerative defect in absence of *Bmal1***

A) Representative Immunoblot of β catenin expression from total and cytoplasmic lung extracts from *Bmal1*$^{creERT2neg}$ *and Bmal1*$^{creERT2/+}$ mice B) quantification of β-catenin expression normalized to β actin from 2 independent experiments (n=4-5) C) Chromatin immunoprecipitation (ChIP) assay of *Bmal1* occupancy in Wnt3a promoter. mPer2 primers were used as positive controls for the analysis. Data are expressed as percentage of input level normalized to IgG control (n=4, pooled from 2 independent experiments).

Tracheal organoids were supplanted with Wnt3a (200ng/ml) and DMSO (0.05% of final concentration) D) *Bmal1*$^{+/+}$ wild type littermates E) *Bmal1*$^{-/-}$ DMSO F) *Bmal1*$^{-/-}$ Wnt3a G) *Bmal1*$^{creERT2neg}$ littermates DMSO H) *Bmal1*$^{creERT2/+}$DMSO I) *Bmal1*$^{creERT2/+}$ Wnt3a, scale bar: 2000µm; CFE for J) *Bmal1*$^{-/-}$ K) *Bmal1*$^{creERT2/+}$. 3 independent experiments with 3 technical replicates. Compiled data are expressed as mean ± SEM. *p<0.01, ****p<0.0001 Kruskal-Wallis test

Scgb1a1$^{Cre/+}$:*Bmal1*$^{fl/fl}$ and their Cre$^{neg}$ littermate were infected as explained in Fig 4E-F and recovered until day 30. Severity of lung injury quantified using the objective histopathological scoring system defined in Fig 1B by a researcher blinded to study group. L-M) Scgb1a1$^{Creneg}$*:Bmal1*$^{fl/fl}$; N-O) Scgb1a1$^{Cre/+}$:*Bmal1*$^{fl/fl}$. Each data point represents individual animal, data pooled from two independent experiments (n=7-8 mice per circadian time point) post 30 days IAV infection. **p<0.005, R; *p<0.01, Mann-Whitney test. $^{\#}$p<0.01, R; $^{\$}$p<0.02, Kruskal-Wallis test.

R) Kaplan-Meier curves for pneumonia separated by quintile of RA score, not controlling for any cofactors.

**Supplemental Figure 1: Gating strategy to isolate lung cells**

A) Gating strategy for isolation of $CD104^{+}$ distal lung airway progenitors and AT2 cells by FACS from *Bmal1*$^{-/-}$, their *Bmal1*$^{+/+}$ wild type littermates, *Bmal1*$^{creERT2/+}$, and *Bmal1*$^{creERT2neg}$. Single cell suspension was sorted for alveolar epithelial cells type II (AT2 as $DARQ7^{-}EPCAM^{+}CD104^{-}$) and $CD104^{+}$basal cells ($DARQ7^{-}EPCAM^{+}CD104^{+}$) post magnetic bead-based depletion of leukocytes and endothelial cells ($CD45^{-}CD31^{-}$).

B) Gating strategy for isolation of lung cells for single cell sequencing from *Bmal1*$^{+/+}$, *Bmal1*$^{-/-}$, *Bmal1*$^{creERT2/+}$, and *Bmal1*$^{creERT2neg}$. Single cell suspension was sorted on $DARQ7^{-}CD31^{-}CD45^{-}$ lung cells for sequencing.

**Supplemental Figure 2: Circadian expression analysis of lung organoids post synchronizing agent treatment.**

Representative records of bioluminescence showing circadian profiles of mPER2 expression in organoids grown from different cellular compartments of lungs from *mPer2*$^{Luc}$ knockin mice treated with dexamethasone (10µM) or Forskolin (100nm) A, D) tracheal organoids (day 7 onwards) B, E) $CD104^{+}$ organoids (day 7 onwards) C, F) AT2 organoids (day 18 onwards)

**Supplemental Figure 3: Circadian clock controls the regeneration of AT2 organoids**

AT2 organoids were raised for following combinations, -A) *Bmal1*$^{+/+}$ AT2 with *Bmal1*$^{+/+}$ fibroblasts B) *Bmal1*$^{-/-}$ AT2 with *Bmal1*$^{+/+}$ fibroblasts C) *Bmal1*$^{+/+}$ AT2 with *Bmal1*$^{-/-}$ fibroblasts D) *Bmal1*$^{-/-}$AT2 with *Bmal1*$^{-/-}$ fibroblasts E) *Bmal1*$^{fl/fl}$*cre*$^{-}$ AT2 with *Bmal1*$^{fl/fl}$*cre*$^{-}$ fibroblasts F) *Bmal1*$^{creERT2/+}$ AT2 with *Bmal1*$^{creERT2neg}$ fibroblasts G) *Bmal1*$^{creERT2neg}$ AT2 with *Bmal1*$^{creERT2/+}$ fibroblasts H) *Bmal1*$^{creERT2/+}$ AT2 with *Bmal1*$^{creERT2/+}$ fibroblasts I) *Bmal1*$^{creERT2neg}$ AT2 with *Bmal1*$^{creERT2neg}$ fibroblasts J) *Bmal1*$^{creERT2neg}$ AT2 with *Cry1,2* DKO fibroblasts K) *Bmal1*$^{creERT2/+}$ AT2 with *Cry1,2* DKO fibroblasts L) Representative Sftpc and Ki67 immunofluorescence images of *Sftpc*$^{CreERT2neg}$*:* Bmal1$^{fl/fl}$ and *Sftpc*$^{CreERt2/+}$*:Bmal1*$^{fl/fl}$ mice explained in Fig 4J-M.

**Supplemental Figure 4: Subtle differences in morphology of organoids from embryonic *Bmal1* knockout**

Representative H&E stained images of embryonic *Bmal1* KO A-B) tracheal organoids, C-D) CD104+ E-F) tracheal organoids and postnatal *Bmal1* KO G-H) CD104+ organoids, scale bar:20µm.

**Supplemental Figure 5: Wnt3a or CHIR did not rescue the *Bmal1* phenotype in AT2 organoids**

Activation of Wnt signaling by either Wnt3a or GSK-3β inhibitor CHIR-99021 in AT2 organoids.

A, E) *Bmal1*$^{creERT2neg}$AT2 with *Bmal1*$^{creERT2neg}$ fibroblasts DMSO B, F) *Bmal1*$^{creERT2neg}$ AT2 with *Bmal1*$^{creERT2neg}$ fibroblasts Wnt3a/CHIR C,G) *Bmal1*$^{creERT2/+}$ AT2 with *Bmal1*$^{creERT2/+}$ fibroblasts DMSO D,H) *Bmal1*$^{creERT2/+}$ AT2 with *Bmal1*$^{creERT2/+}$ fibroblasts Wnt3a/CHIR respectively, I)CFE. 3 independent experiments with 3 technical replicates, scale bar: 2000µm. Compiled data are expressed as mean ± SEM. ****p<0.0001.
