## Supplementary material for "Circadian regulation of lung repair and regeneration": Table 1

Table 2: Primers used in ChIP experiment

| Primer | Forward | Reverse |
|---|---|---|
| mPer2 | GGTTCCGCCCCGCCAGTATGC | CCGTCACTTGGTGCGCTCGGC |
| *Bmal1*_Wnt3a location1 | TCCTGCTGAGCTTCCTTTAC | CCTGAGTCCATTTGCAGAATTT |
| *Bmal1*_Wnt3a location2 | GGACTACGCAGTGCAATTCT | GTGTGTCGGTGTTCGTAGAG |
