## Supplementary material for "Circadian regulation of lung repair and regeneration": Table 2

Table 2: Period and phase of PER2::LUC lung organoids in presence and absence of synchronizing agents

| Type of organoid | Synchronizing agent | Period (h) | Phase (h) | Amplitude |
|---|---|---|---|---|
| Tracheal | - | 26-28.5 | 0.5-1.8 | 75825-116050 |
| | Dexamethasone | 27.1-27.6 | 9.7-10.3 | 785717-1261026 |
| | Forskolin | 21.5-28.9 | 0.6-10.7 | 199481-487321 |
| CD104+ | - | 24.3-26.3 | 5.0-9.4 | 10131-45178 |
| | Dexamethasone | 25.8-26.5 | 9.4-9.9 | 15543-39601 |
| | Forskolin | 26.9-27.2 | 9.9-10.4 | 5368-7343 |
| AT2 | - | 26.9-27.5 | 7.0-9.0 | 18860-47285 |
| | Dexamethasone | 26.8-27.2 | 9.6-10.5 | 209273-778301 |
| | Forskolin | 27.7-29.2 | 8.1-9.0 | 150721-284728 |
