## Supplementary material for "Circadian regulation of lung repair and regeneration": Table 3

Table 3: Period and phase of Wnt/ β-catenin targets (http://circadb.hogeneschlab.org/mouse)

| Gene Name | Period (h) | Phase | Q value (JTK) |
|---|---|---|---|
| **Axin2** | 24 | 19 | 0.00110893 |
| Ccnd1 | 26 | 7 | 0.000255309 |
| Scd2 | 24 | 17 | 0.0004672 |
| Wee1 | 24 | 13 | 9.20767e-09 |
